# Multi-colour smFISH with automated cell-cycle quantification reveals distinct localization of co-regulated cell wall mRNAs in *Saccharomyces cerevisiae* yeast

**DOI:** 10.64898/2026.11.17.763897

**Authors:** Marah M. Jnied, Sander C. van Otterdijk, Kelly van Rossum, Florence Berbain, Carlotta Meyer, Philipp Savakis, Evelina Tutucci, Edward W. J. Wallace

**Affiliations:** Institite for Cell Biology and Centre for Engineering Biology, School of Biological Sciences, University of Edinburgh; Systems Biology Section, A-LIFE Department, Amsterdam Institute of Molecular and Life Sciences (AIMMS), Vrije Universiteit Amsterdam, Amsterdam, the Netherlands

## Abstract

Every aspect of messenger RNA life and fate is dictated by its complementary bound proteins. RNA-binding proteins (RBPs) regulate mRNA localization, translation, and/or degradation, and mRNAs encoding functionally related proteins can form “RNA regulons” that are bound by shared sets of RBPs. Here, we present an open-source multi-colour smFISH-based workflow we developed to systematically quantify mRNA abundance and localization across the budding yeast cell cycle. With this workflow, we investigate the cell-cycle dependent regulation of mRNAs encoding cell wall proteins (cwmRNAs) in *Saccharomyces cerevisiae* yeast. We quantify the localization of *SRL1* and *SUN4* cwmRNAs which are bound by Ssd1, Mpt5/Puf5, and She2-She3 among other RBPs. We show that *SRL1* and *SUN4* exhibit distinct cell cycle-dependent abundance and localization patterns, and that *SRL1* mRNA relies on She2 and She3 and partially Ssd1, but not Mpt5, for its localization to the bud. Our results argue that Ssd1 and Mpt5 are not primarily RNA localization factors and confirm the importance of the She2-She3 complex in conferring bud-localization to diverse mRNAs. We also show that *SUN4* and *SRL1* mRNA localization is not affected by the Sec complex which facilitates the localization of their protein products. These findings show that co-regulated mRNAs can follow distinct spatiotemporal programs, and our workflow provides a general framework to test how RNA-binding proteins coordinate mRNA localization in single cells.

## Introduction

The fate of a messenger RNA – where it is localized, when and how efficiently it is translated, and how long it persists – is determined by the RNA-binding proteins (RBPs) that associate with it (Singh et al. 2015). mRNAs encoding functionally related proteins are often bound by shared sets of RBPs, which have been proposed to act as post-transcriptional “RNA regulons” that coordinate gene expression, by analogy with bacterial operons (Keene 2007). Systematic studies of RBPs in *Saccharomyces cerevisiae* support the RNA regulon concept. For example, Puf-family proteins bind mRNAs encoding proteins of the same subcellular compartment or pathway, such as mitochondrial proteins (D. J. Hogan et al. 2008; G. J. Hogan et al. 2015), and the She2–She3 complex binds and transports a set of mRNAs to the bud, including *ASH1* and *CLB2* (Böhl et al. 2000; Long et al. 2000; Shepard et al. 2003; Maekiniemi et al. 2025; Das et al. 2021; van Otterdijk et al. 2026). mRNAs encoding cell wall proteins (cwmRNAs) are another example: they are co-bound by Ssd1, She2, Khd1/Hek2, Mpt5/Puf5, Bfr1 and Scp160 (D. J. Hogan et al. 2008; Hall and Wallace 2022). Shared RBPs could coordinate gene expression in space, by localizing mRNAs to a subcellular site, and in time, by acting at particular phases of the cell cycle through transcription, processing, translation or decay. Whether co-bound mRNAs are in fact found at the same place and time within individual cells, however, remains largely untested.

RNA regulons have mostly been defined by bulk methods, such as RBP immunoprecipitation followed by microarrays or sequencing, crosslinking-based approaches such as CRAC, and RNA tagging (D. J. Hogan et al. 2008; Lapointe et al. 2017; Miller et al. 2018; Bayne et al. 2021). Cell cycle-dependent expression has similarly been measured across synchronized populations (Spellman et al. 1998; Kelliher et al. 2016; Eser et al. 2014). These approaches do not detect where each mRNA is within the cell, nor how mRNA abundance varies between individual cells. Synchronization can also perturb the cell cycle that it aims to resolve. As a result, the spatial and temporal behaviour of most regulon members, and how their shared RBPs control it, has been inferred rather than measured.

Single-molecule fluorescence in situ hybridization (smFISH) counts and localizes individual endogenous mRNAs in fixed cells by microscopy and image analysis (Femino et al. 2003; Raj et al. 2008; Zenklusen et al. 2008). Multi-colour smFISH can detect several mRNAs in the same cell, and nuclear staining and cell morphology simultaneously report cell-cycle phase (Tutucci and Singer 2020; Maekiniemi et al. 2022). Spot detection (Mueller et al. 2013; Imbert et al. 2021) and deep learning-based cell segmentation (Stringer et al. 2021; Cutler et al. 2022) are now largely automated. In budding yeast, however, measuring cell-cycle dependent localization also requires identifying the bud, linking it to its mother cell and assigning a cell-cycle phase. These steps are usually done by hand, which limits throughput and reproducibility. No reliable, automated tools currently exist to quantify mRNA localization to the bud across the cell cycle.

Here, we developed an smFISH-based workflow to quantify mRNA localization to the bud across the yeast cell cycle. *CLB2* and *ASH1* mRNAs are co-labelled in a single channel as a bud marker, which leaves two channels free for mRNAs of interest (Maekiniemi et al. 2022, 2025). We present two open-source Python packages, buddy and napari-buddy, that combine cell segmentation, spot detection, correction of channel offsets and assisted mother–bud assignment, and assign cell-cycle phase automatically; we validated these assignments against blinded manual annotation.

We applied this workflow to two cwmRNAs, *SRL1* and *SUN4*, which are bound by Ssd1 (D. J. Hogan et al. 2008; Jansen et al. 2009; Bayne et al. 2021) and also by Mpt5/Puf5 (Lapointe et al. 2017; Miller et al. 2018). We then tested the contribution of factors previously implicated in their regulation. Ssd1 is required for yeast growth, aneuploidy tolerance, longevity, thermotolerance and resistance to cell wall stress (Uesono et al. 1997; Kaeberlein and Guarente 2002; Mir et al. 2009; Hose et al. 2020; Breeden and Miles 2022). Ssd1 has been proposed to repress translation of cwmRNAs (Jansen et al. 2009; Kleinerman et al. 2026) and to promote bud localization of *SRL1* mRNA (Kurischko et al. 2011). Mpt5/Puf5 is required for cell wall integrity and regulates *CLB2* expression (Kaeberlein and Guarente 2002; Stewart et al. 2007; Sato et al. 2025). The She2–She3 complex tethers mRNAs to the Myo4 motor for transport to the bud, and has been reported to bind *SRL1*, and possibly *SUN4*, mRNAs (Böhl et al. 2000; Long et al. 2000; Shepard et al. 2003; Oeffinger et al. 2007). Finally, the Sec complex subunit Sec72 is required to target Srl1 and Sun4 proteins to the ER (Ast et al. 2013), allowing us to ask whether disrupting protein targeting affects mRNA localization.

We find that *SRL1* and *SUN4* mRNAs, despite sharing RBPs, differ in when and where they are found. *SRL1* mRNA peaks in S and G2 phase and is strongly localized to the bud; its bud localization requires She2 and She3 and is partially reduced in the absence of Ssd1. In contrast, *SUN4* mRNA shows a different expression pattern and is not enriched in the bud. Mpt5 affects the abundance of *CLB2* and *SUN4* mRNAs but not their localization, and Sec72 affects neither mRNA’s localization or abundance. Together, these results show that shared RBP binding does not imply shared localization, and that members of a proposed RNA regulon can be regulated with distinct spatiotemporal patterns.

## Results

### A combined *CLB2* + *ASH1* smFISH signal marks the bud throughout the budded cell cycle

To quantify mRNA localization to the bud, we first needed a marker that identifies the bud in fixed cells throughout the cell cycle. For this we used two well-characterized bud-localized mRNAs with complementary expression windows (Figure 1A–B) (Tutucci et al. 2018; Maekiniemi et al. 2022, 2025). *CLB2* mRNA is expressed from S phase through early mitosis, and *ASH1* mRNA accumulates in anaphase. Both are transported to the bud by the She2–She3 complex (Böhl et al. 2000; Long et al. 2000; Takizawa and Vale 2000; Maekiniemi et al. 2025). Labelling both mRNAs with probes carrying the same fluorophore therefore gives a single bud marker from bud emergence to cytokinesis and leaves the remaining two channels of a three-colour smFISH experiment free for two mRNAs of interest (Figure 1C, panel i). Because *CLB2* and *ASH1* are not expressed in G1, the combined signal also helps distinguish unbudded G1 cells from budded cells, which we used for cell-cycle assignment (see below).

**Figure 1.**
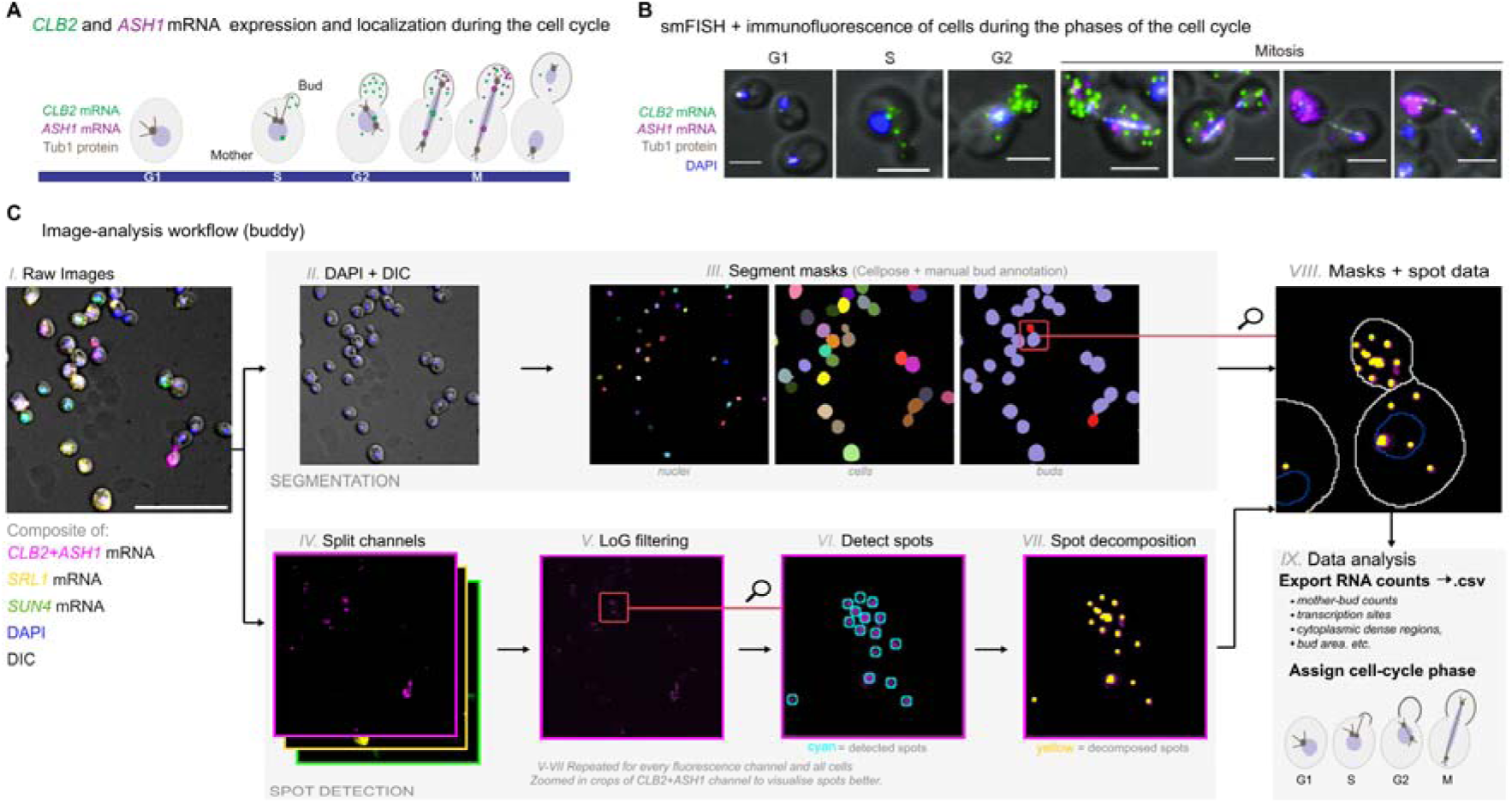
Co-labeling of *CLB2* and *ASH1* mRNAs marks the bud throughout the cell cycle, enabling automated quantification of mRNA localization. A, Schematic of *CLB2* and *ASH1* mRNAs expression during the cell cycle. Green dots represent *CLB2* mRNAs. Magenta dots represent *ASH1* mRNAs. Tub1 (grey) co-localizes with the spindle pole body, which is duplicated during S phase. Bud emergence starts during S phase and ends with formation of the daughter cell. During anaphase, the Tub1-marked microtubules stretch between the mother and the daughter cell. **B**, Merged maximal z-projections of *CLB2* mRNA smFISH (green), *ASH1* mRNA smFISH (magenta), anti-Tub1 IF (white) and DAPI (blue) merged to a single DIC section (grey). The corresponding cell cycle phase is indicated on top of the panel. Scale bars 3 μm. **C**, Image-analysis workflow implemented in buddy and napari-buddy. (i) Input data, shown as a composite image of DIC (grey) and fluorescence channels: *CLB2*+*ASH1* mRNA (magenta), *SRL1* mRNA (yellow), *SUN4* mRNA (green) and DAPI (blue). Scale bar, 20 µm. (ii) The DAPI and DIC channels are extracted and (iii) used to segment nuclear and cell masks with Omnipose. Bud masks are used to manually link buds to their mother cells in napari-buddy. (iv) Each fluorescence channel is split and processed for spot quantification: (v) Gaussian background subtraction, (vi) spot detection by Laplacian-of-Gaussian filtering and local-maximum thresholding, and (vii) decomposition of dense spot regions with Big-FISH. Panels vi – vii show a zoomed crop of the *CLB2*+*ASH1* channel for clarity; the analysis was performed on every cell in the image. (viii) Masks and spot data are aligned and combined to assign RNAs to mother and bud compartments. (ix) Per-cell features, including mother and bud RNA counts, transcription sites, cytoplasmic dense regions and bud area, are exported as a.csv file and used to assign each cell automatically to a cell-cycle phase based on bud area, the number and position of nuclei, and *CLB2*+*ASH1* mRNA counts (see Methods). Panels A and B are adapted from our preprint (Maekiniemi et al. 2022); these data were not included in the peer-reviewed version of that study (Maekiniemi et al. 2025).

### Semi-automated quantification of mRNA localization across the cell cycle with buddy and napari-buddy

To analyse these images systematically, we developed two open-source Python packages: buddy (https://github.com/Tutuccilab/buddy) and its napari plugin, napari-buddy (https://github.com/Tutuccilab/napari-buddy). They combine established tools for cell segmentation (Omnipose; (Cutler et al. 2022)), spot detection (Big-FISH; (Imbert et al. 2021)) and image visualization (napari; (Chiu et al. 2022)) into a single workflow with five steps (Figure 1C):

1. **Segmentation.** Omnipose segments nuclei and cells from the DAPI and DIC channels (panels ii–iii).
2. **Spot detection.** In each fluorescence channel, Big-FISH detects individual mRNAs and estimates how many molecules are in dense regions such as transcription sites (panels iv–vii).
3. **Automated set-up steps.** buddy automatically selects in-focus z-slices and corrects the offset between the DIC and fluorescence channels. Detection parameters are chosen interactively in napari-buddy on a representative image, then saved and reused to batch-process all images of an experiment.
4. **Manual curation.** The only manual steps are correcting segmentation masks where needed and linking each bud to its mother cell, which napari-buddy speeds up by letting the user draw these links directly on the image.
5. **Output.** mRNAs are assigned to the mother or bud compartment (panel viii), and per-cell features are exported for analysis (panel ix): total mRNA count, mother and bud counts, transcription sites, bud area and nuclear position.

From these per-cell features, cells are assigned to a cell-cycle phase (Methods; Figure 1C, panel ix). G1 cells are unbudded and contain five or fewer *CLB2*+*ASH1* mRNAs; S- and G2-phase cells have a single nucleus in the mother and a bud area below or above 2 µm², respectively; and M-phase cells have a nucleus in the bud. To validate this assignment, four individual human annotators each classified a random subset of cells from all strains without seeing the automatic call. Automatic and manual calls agreed for 84.7% of annotations (Cohen’s κ = 0.80). Agreement was highest for G1 and M phase (precision 94% and 90%) and lowest for S phase (72%), which was mostly confused with the neighbouring G1 or G2 phases (Figure S1).

For each mRNA we report two measures per cell: the total number of mRNAs, and the proportion of those mRNAs located in the bud. Across 7 strains and 3–5 biological replicates per strain, we analysed 12,526 cells, of which 2,406 (19%) were budded.

Having established this workflow, we applied it to two cell wall mRNAs, labelled in separate fluorescent channels alongside the *CLB2* + *ASH1* bud marker.

### *SRL1* and *SUN4* mRNAs have distinct abundance and localization patterns over the cell cycle

We selected 2 cwmRNAs that are prominent targets of both Ssd1 and Mpt5: *SRL1* and *SUN4* (D. J. Hogan et al. 2008; Jansen et al. 2009). Both these are over 100-fold enriched in Ssd1 binding and contain Ssd1-binding sites proximal to the start codon, as measured by cross-linking and analysis of cDNA (CRAC) (Bayne et al. 2021). Both are also bound by Mpt5/Puf5 as shown by independent sequencing assays, based on RNA immunoprecipitation by tagged Mpt5 (RIP-seq) (Miller et al. 2018) or RNA modification by a Poly(U)-polymerase enzyme fused to Mpt5 (Lapointe et al. 2017). *SRL1* encodes a cell wall mannoprotein that is required to compensate for cell wall instability (Hagen et al. 2004), and Srl1 protein is localized to the bud (Shepard et al. 2003; Kurischko et al. 2011). *SRL1* mRNA localization was previously claimed to depend on Ssd1 (Kurischko et al. 2011), and *SRL1* mRNA is bound by She2 (Shepard et al. 2003; Oeffinger et al. 2007; D. J. Hogan et al. 2008). Indeed, a fragment of *SRL1* mRNA confers both She2-She3 binding and bud-localization (Jambhekar et al. 2005). *SUN4* encodes the Sun4/Scw3 protein whose absence results in defective mother-daughter cell separation in *S. cerevisiae* (Mouassite et al. 2000), and is part of the “SUN family” of β-glucan binding proteins that have a conserved role in fungal cell wall biogenesis and septation (Firon et al. 2007; Gastebois et al. 2013; Schöppner et al. 2026). *SUN4* mRNA may also be bound by She2 (Oeffinger et al. 2007). We recently found that Ssd1 represses Sun4 protein abundance post-transcriptionally, by binding the *SUN4* 5′ UTR (Kleinerman et al. 2026). Sun4 protein localizes to the ER, bud neck, cell septum, and bud scar (Kuznetsov et al. 2016; Dubreuil et al. 2019), but *SUN4* mRNA localization was not previously tested.

Our multi-colour smFISH workflow reveals patterns of abundance and localization of *SRL1* and *SUN4* mRNAs during the yeast cell cycle (Figure 2). *SRL1* becomes abundant during S-phase during bud emergence (Figure 2A, 2B) and localizes to the bud throughout S to M-phase (Figure 2A, 2C). As expected, *CLB2* + *ASH1* mRNAs localize to the bud, similarly to *SRL1* mRNA, and are more abundant in G2 and M-phase. By contrast, *SUN4* mRNA is slightly more abundant during G1 compared to S phase (Figure 2B) and does not clearly localize to the bud (Figure 2A, 2C). For each mRNA, proportion of mRNA in the bud across all budded phases is consistent across replicates (Figure 2D).

**Figure 2:**
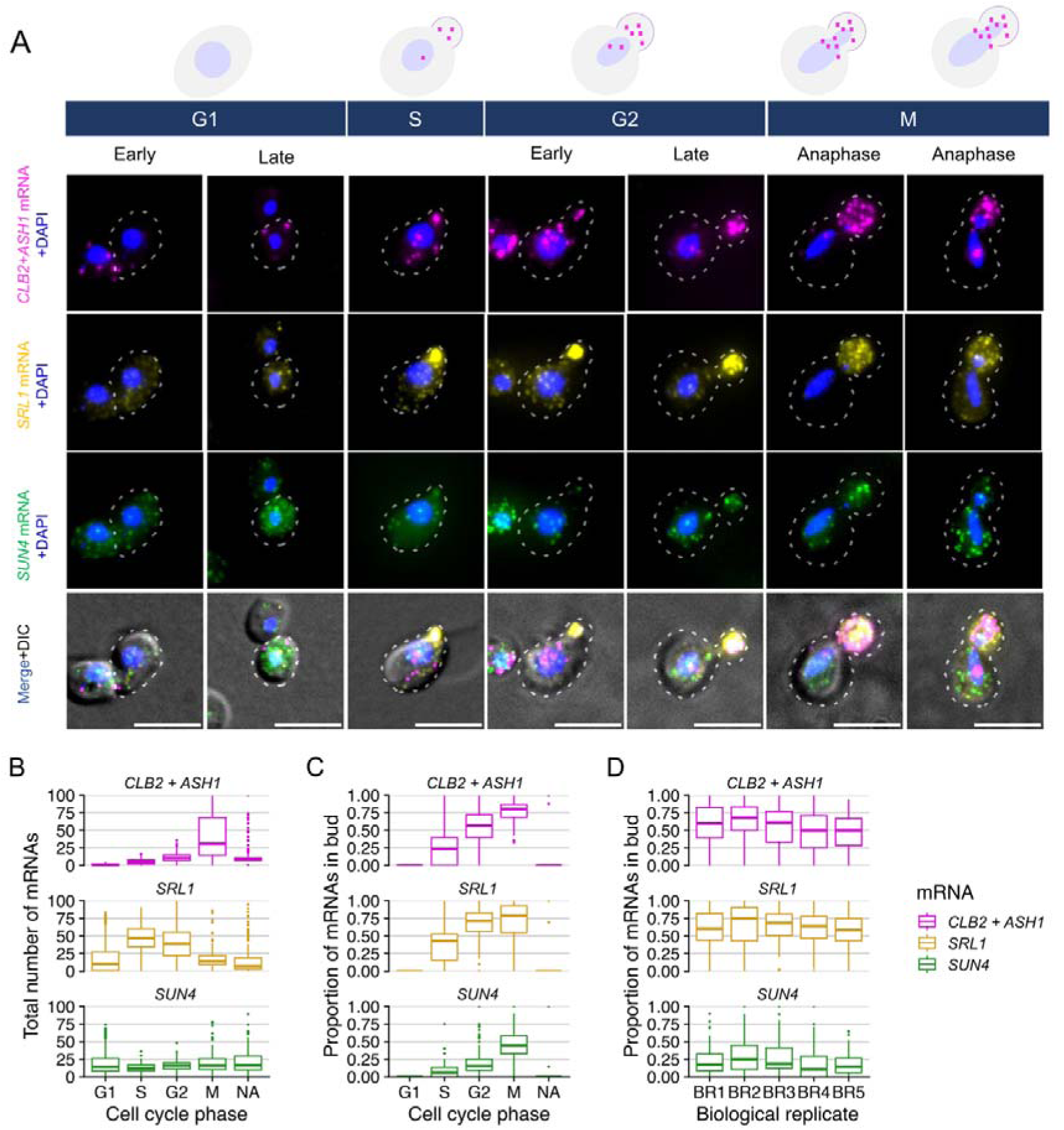
***SRL1* and *SUN4* mRNAs have distinct abundance and localization patterns over the cell cycle. A,** Top, schematic of the budding yeast cell cycle and mRNA bud-localization. Bottom, Images of representative cells (wild-type BY4741) from phases of the cell cycle (G1, S, G2, and M); from top to bottom, smFISH images show merged maximal z-projections of *CLB2 + ASH1* mRNA (magenta), *SRL1* mRNA (yellow), *SUN4* mRNA (green), and merged images with all three mRNAs and DIC (gray) for cell outlines. All images show the cell outline in dashed white line, and the nucleus stained by DAPI (blue). Scale bars are 5 μm. **B,** Total mRNA counts per cell, for each assigned cell cycle phase. Counts of *CLB2 + ASH1* mRNAs increase through S, G2, and M phases, while counts of *SRL1* mRNA are highest in S and G2 phase, and counts of *SUN4* mRNA are slightly lower in S phase. **C**, Proportion of mRNA in bud per cell for each assigned cell cycle phase; this is non-zero for budded cells which are found only in S, G2, and M phases. Positive control *CLB2 + ASH1* mRNAs are bud-localized. *SRL1* mRNA is strongly bud-localized with the median cell having 43% of mRNAs in the bud in S-phase and 71% in the bud in G2-phase. By contrast, *SUN4* mRNA is not clearly bud-localized with the median cell having 6% of mRNAs in the bud in S-phase and 15% in the bud in G2-phase. In the boxplots in panels B and C, results are pooled across 5 biological replicates (n = 1,696 cells: G1, 1,266; S, 95; G2, 213; M, 122). 188 cells were not assigned to a cell cycle phase (NA) are shown. **D,** Proportion of mRNA in bud per cell across all budded cells, regardless of cell cycle phase assignment, for each of 5 biological replicates (n = 42, 132, 117, 61 and 87 budded cells; 439 in total).

### *SRL1* mRNA relies on She2, She3 and partially Ssd1 for its localization to the bud

We next asked how the RNA-binding proteins Ssd1, Mpt5, She2, and She3 contribute to the abundance and localization of *SRL1* and *SUN4* mRNAs, by applying the same multi-colour smFISH and data analysis approach as above to deletion strains (Figure 3, S3, S4, S5).

**Figure 3:**
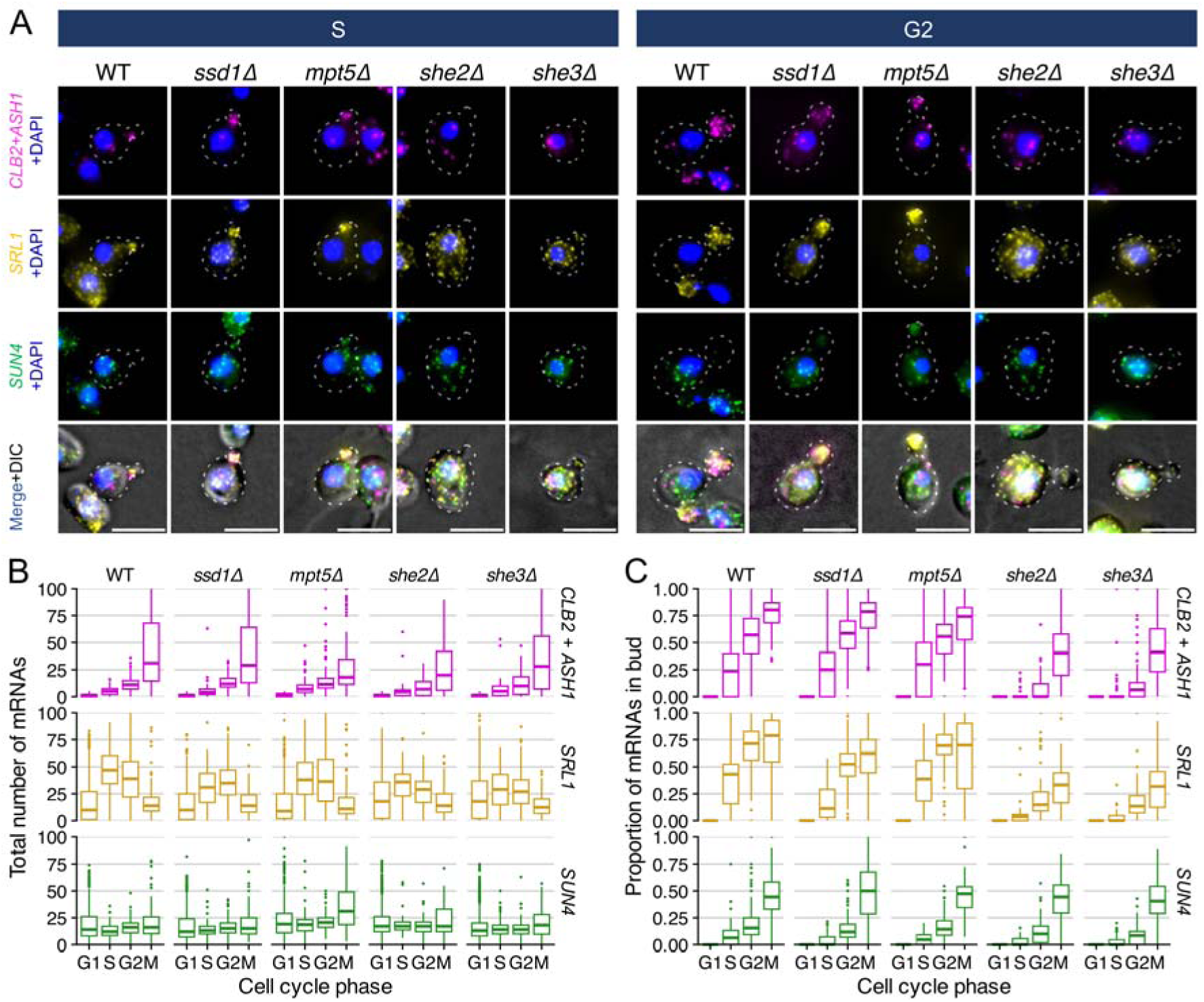
***SRL1* mRNA relies on She2, She3 and partially Ssd1 for its localization to the bud. A,** Images of representative cells from S phase and G2 phase of cell cycle, for wild-type (BY4741), *ssd1*Δ, *mpt5*Δ, *she2*Δ and *she3*Δ cells. From top to bottom, smFISH images show merged maximal z-projections of *CLB2 + ASH1* mRNA (magenta), *SRL1* mRNA (yellow), *SUN4* mRNA (green), and merged images with all three mRNAs and DIC (gray) for cell outlines. Scale bars are 5 μm. **B,** Total mRNA counts per cell for each assigned cell cycle phase across these strains. **C,** Proportion of mRNA in bud per cell for each assigned cell cycle phase across these strains. While *SRL1* mRNA is less bud-localized in *ssd1*Δ or *mpt5*Δ compared to wild-type, its bud-localization is abolished in *she2*Δ and *she3*Δ cells. In the boxplots in panels B and C, results are pooled across 3 to 5 biological replicates (number of cells in G1 / S / G2 / M: wild-type, 1,266 / 95 / 213 / 122 (5 replicates); ssd1Δ strain, 1,567 / 73 / 235 / 117 (4); mpt5Δ strain, 759 / 64 / 154 / 71 (3); she2Δ strain, 1,207 / 56 / 127 / 69 (3); she3Δ strain, 1,343 / 81 / 163 / 72 (5)), and the wild-type data shown is the same as in Figure 2B and 2C. Matched data from G1 and M phase are shown in Figures S3 and S4.

In *ssd1*Δ cells, we observed that for control *CLB2 + ASH1* mRNA the overall patterns of mRNA abundance and localization are similar to wild-type (Figure 3A-C). *SRL1* mRNA has a peak of abundance during S-phase in wild-type cells, which is decreased about 1/3 in *ssd1*Δ cells (Figure 3B). Additionally, *SRL1* mRNA is less bud-localized in *ssd1*Δ than in wild-type cells, for example the median S-phase cell has 43% of *SRL1* transcripts bud-localized in wild-type but only 11% in *ssd1*Δ. This partial mis-localization extends to G2-phase, in which the median cell has 71% of *SRL1* transcripts bud-localized in wild-type and only 52% in *ssd1*Δ. Lower bud localization of *SRL1* mRNA in *ssd1*Δ compared to wild-type is consistent across biological replicates, and is statistically significant according to t-test on the difference in mean proportion in bud (Figure S5). Meanwhile, there are no substantial changes in abundance or localization of *SUN4* mRNA in *ssd1*Δ cells compared to wild-type, although the slightly lower bud-localization of *SUN4* mRNA in S-phase *ssd1*Δ cells is still scored as statistically significant (Figure S5).

In *mpt5*Δ cells, abundance of *CLB2 + ASH1* mRNA is decreased in M-phase, accompanied by a slight decrease in bud-localization. In the median M-phase cell, abundance of *CLB2 + ASH1* mRNA decreases to around 13 mRNAs per cell in *mpt5*Δ cells, in comparison to 30 in wild-type. This is consistent with previous studies showing that Mpt5p positively regulates the cell cycle specific expression of *CLB2* and that *CLB2* mRNA levels are reduced in *mpt5*Δ strains (Sato et al. 2025). In contrast, *SUN4* total mRNA abundance in *mpt5*Δ is slightly higher than wild-type cells in S to M-phase. However, bud-localization of *CLB2 + ASH1* mRNA or of *SRL1* mRNA is unperturbed in S-phase or G1-phase *mpt5*Δ cells.

In *she2*Δ and *she3*Δ cells, *SRL1* mRNA localization to the bud is abolished (Figure 3). The median S-phase cell has 5% of *SRL1* localized to the bud in *she2*Δ and 0% in *she3*Δ, in comparison to 43% in wild-type. The median G2-phase cell has 15% of *SRL1* localized to the bud in *she2*Δ and 14% in *she3*Δ, in comparison to 71% in wild-type. These differences are consistent across biological replicates and the difference in means is statistically significant (p < 0.001; Figure S5). As expected, similar patterns of abolished bud-localization can be seen in control *CLB2 + ASH1* mRNA in *she* mutant strains. We observed *SUN4* mRNA bud-localization is even lower in she2Δ and she3Δ than in wild-type (Figure S5). Abundance of *CLB2 + ASH1* mRNA is perhaps slightly decreased in *she* mutant strains. Meanwhile, *SUN4* mRNA abundance is not substantially affected.

### *SRL1* mRNA bud localization is quantitatively enriched relative to bud size

We sought a continuous quantitative analysis of mRNA abundance and localization relative to cell size and bud size in our smFISH dataset, to complement the above analysis where cells were binned into cell-cycle phases. We use cross-sectional area from DIC single focal z-plane images as our proxy for cell size, abbreviated as “area”.

Firstly, we plotted total mRNA abundance against cell size, summarized with a smoothed mean of mRNA abundance per cell plotted using a generalized additive model (Figure 4A). For *CLB2 + ASH1* mRNA in wild-type, small cells with area less than 15 µm^2^ have less than 5 mRNAs per cell on average. As cells grow beyond 15 µm^2^, the mean abundance rises and peaks above 35 mRNAs per cell for the largest cells. This pattern is similar in *ssd1*Δ and *mpt5*Δ, but in *she2*Δ and *she3*Δ strains larger cells have less than 20 mRNAs per cells. The 95% confidence intervals from the smoothed fit are very small (shading around trendline), indicating that these differences in mean abundance are statistically significant despite large cell-to-cell variability.

**Figure 4.**
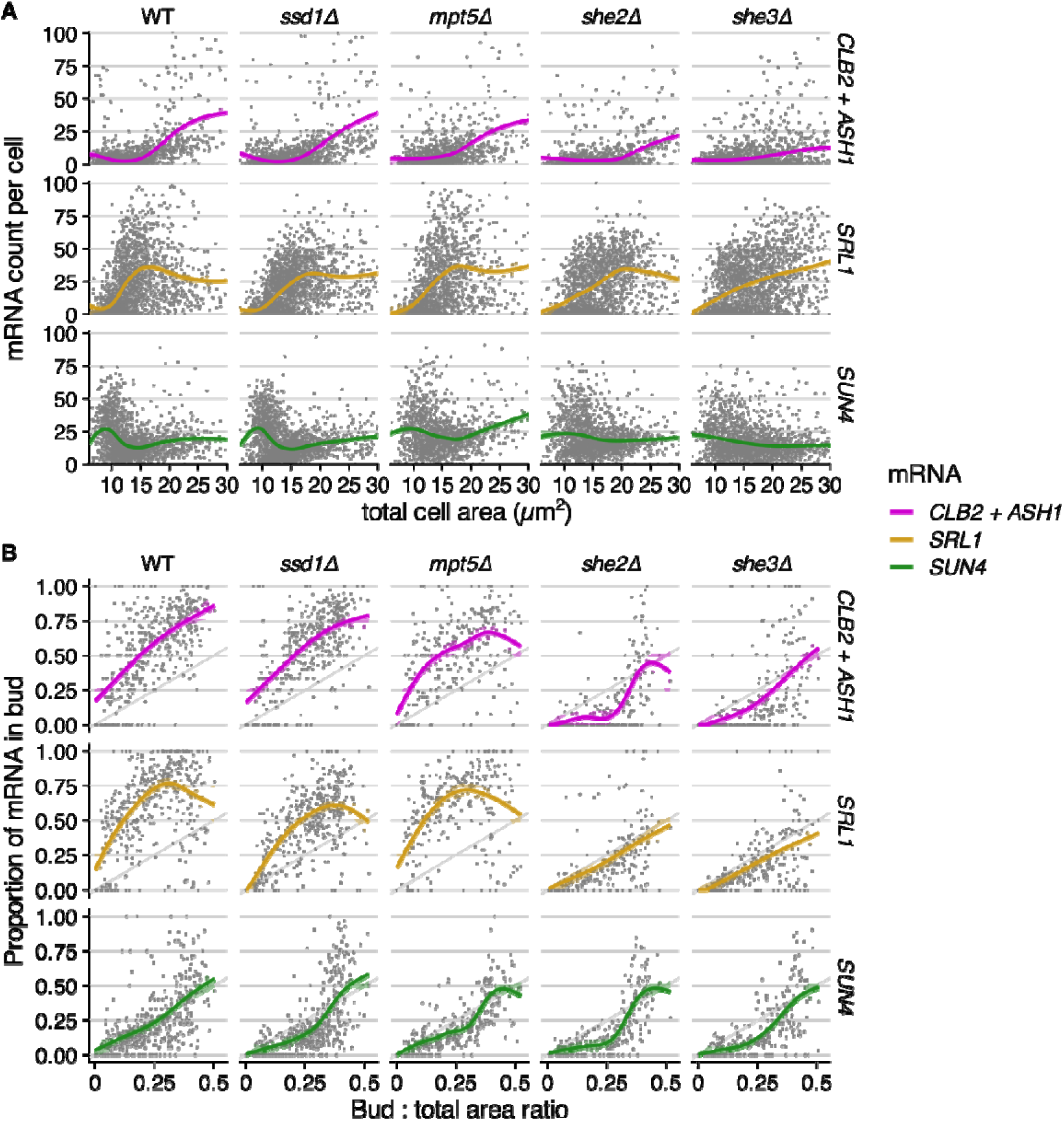
RNA abundance and localization vary smoothly depending on cell size and bud:total size ratio. A, Total mRNA abundance against cell size (total cross-sectional area in µm^2^), for all strains. **B,** Proportion of mRNA in bud plotted against bud:total area ratio, for budded cells in all strains. In all panels the trendline summarizes the y-axis as a smoothed mean calculated using a generalized additive model applied to the x-axis (area), and shading around the trendline shows a 95% confidence interval, i.e. smoothed standard error. The underlying data are the same as in Figure 3B and 3C, and results are pooled across 3 to 5 biological replicates per strain.

Mean *SRL1* mRNA abundance per cell is around 5 in the smallest wild-type cells but rises rapidly to a peak of 35 mRNAs per cell in cells with area of 15 µm^2^. For even larger cells, *SRL1* mRNA abundance per cell declines to 25 mRNAs per cell. These patterns are qualitatively similar in *mpt5*Δ*, ssd1*Δ, *she2*Δ, and *she3*Δ strains, but shifted so that peak abundance occurs in larger cells relative to wild-type.

*SUN4* mRNA abundance peaks in small cells with an area of less than 10 µm^2^, at about 25 mRNAs per cell in wild-type and *ssd1*Δ strains. The abundance then declines to a mean of 12 mRNAs per cell for cells with an area of 15 µm^2^, slightly increasing for larger cells. Size-dependent variations in *SUN4* mRNA abundance are subtly different in *mpt5*Δ strains, and attenuated in *she2*Δ and *she3*Δ strains.

We similarly analyzed nascent mRNA counts in the mother cell nucleus compared to total cell area (Figure S6), which show trends consistent with total mRNA. For example, in wild-type cells mean nascent *CLB2 + ASH1* mRNA increases in cells with total area over 15 µm^2^, mean nascent *SRL1* mRNA is significantly above zero only for cells with area 10 to 15 µm^2^, and mean nascent *SUN4* mRNA peaks for cells with area under 10 µm^2^.

Next, we estimated mRNA localization in the bud at each cell size by comparing bud:total mRNA counts with bud:total area ratio (Figure 4B). In the null expectation of no bud localization, the mRNA count ratio would track the area ratio. In wild-type cells, *CLB2 + ASH1* mRNAs are enriched in the bud, with 18% of mRNAs in the bud on average as soon as the bud can be detected, and 80% of mRNAs in the bud when the bud reaches 50% of the total cell area. Again, 95% confidence intervals calculated from the smoothed fit are very small (shaded area around magenta trendline), indicating bud localization is statistically significantly above the null expectation (grey diagonal line). *SRL1* mRNA is also strongly enriched at the bud, especially for intermediate-sized buds where about 75% of *SRL1* mRNA are in the bud when the bud is 30% of the total cell area. *SUN4* mRNA resembles the null expectation, with proportion of mRNA in bud closely tracking the bud:total area ratio, such that confidence intervals overlap the diagonal line. While *ssd1*Δ and *mpt5*Δ have small effects on bud localization of mRNA, *she2*Δ and *she3*Δ abolish it. Bud localization of *SRL1* mRNA decreases to below null expectations in both *she2*Δ and *she3*Δ strains.

This quantitative analysis confirms that *SUN4* and *SRL1* mRNAs have distinct cell-size dependent expression, and that *SRL1* is strongly bud-localized dependent on the She complex with a contribution from Ssd1. *SUN4* mRNA abundance peaks in smaller cells compared to *SRL1* mRNA abundance, and average expression of *SRL1* and *SUN4* mRNAs are anti-correlated. In wild-type cells, *SRL1* mRNA localizes to the bud at all bud sizes, unlike *SUN4* mRNA. Cell-size-dependent mRNA abundance and localization is barely affected in *ssd1*Δ or *mpt5*Δ strains. Disrupting the She2-She3 complex perturbs cell-size-dependent mRNA abundance and abolishes localization of *CLB2*, *ASH1*, and *SRL1* mRNAs to the bud.

### *SUN4* and *SRL1* mRNA localization does not depend on the Sec complex

Cell wall proteins with N-terminal cleaved signal peptides, such as *SUN4* and *SRL1*, depend on the Sec complex to translocate into the ER as the first step of their secretion to the cell wall (Ast et al. 2013). In strains with mutations to the Sec complex, tagged cell wall proteins including Sun4 and Srl1 are mislocalized to cytoplasmic aggregates (Ast et al. 2013). We asked if the Sec complex mutant sec72Δ also affected localization of *SUN4* and *SRL1* mRNAs. The smFISH data showed no significant changes in mRNA localization compared to wild-type (Figure S7), indicating that mRNA localization does not depend on Sec72.

## Discussion

We developed an experimental and computational workflow to quantify abundance and localization of mRNAs across the budding yeast cell cycle (Figure 1), and applied it to investigate cell wall mRNAs as a model RNA regulon. Multi-colour smFISH enabled detection of two cwmRNAs in the same cells as bud marker mRNAs, so that differences in localization could be compared directly within a cell rather than inferred across separate populations. Combining smFISH with DAPI staining of the nucleus and DIC imaging of the cell outline allowed the cell-cycle phase of individual cells to be determined. This process was semi-automated by our buddy pipeline, with manual input limited to correcting segmentation masks and linking mother–bud pairs, the latter sped up by a custom napari plugin, napari-buddy. Automated correction of the offset between DIC and fluorescence channels, and validation of cell-cycle calls against blinded manual annotation, made this analysis scalable across multiple strains and biological replicates. The remaining manual step, linking buds to mothers, can now be addressed by a U-Net-based bud segmentation model trained on this dataset (Figure S2).

Our workflow enabled two kinds of quantitative analysis; either categorizing cells by cell cycle phase (Figure 2, 3) or, without explicitly assigning phase, analyzing mRNA abundance and localization dependent on cell size and bud size (Figure 4). The results are consistent and complementary, with cell-cycle phase analysis clearly indicating that *SRL1* mRNA localizes to the bud at all budded phases. A remaining limitation in the cell cycle assignment is accurately identifying S phase, defined here by small bud size; additional cell-cycle markers or live-cell imaging could refine these calls. Complementing this, the size-dependent analysis enabled a more precise assessment of mRNA regulation, showing that *SUN4* mRNAs peak in average abundance in smaller cells than *SRL1* mRNAs. Strikingly, *ASH1* + *CLB2* mRNAs and *SRL1* mRNA are quantitatively enriched in the bud (Figure 4). However, a non-localized *SUN4* mRNA is detected in the bud proportional to the ratio of bud size to total size, matching the null expectation from a spatially uniform distribution. This combination of multi-colour smFISH with semi-automated single-cell analysis will be broadly applicable to study co-regulation of other mRNAs in heterogenous cell populations.

Our analysis revealed that cell wall mRNAs (cwmRNAs) have diverse spatiotemporal distributions over the yeast cell cycle, despite being bound by overlapping sets of RNA-binding proteins (RBPs) in a proposed RNA regulon. *SRL1* mRNA is localized to the yeast bud but *SUN4* mRNA is not, despite both being bound by Ssd1, Mpt5, and other shared RNA-binding proteins (Figure 5). This argues that the cell wall RNA regulon operates in a more nuanced way than Keene’s proposed analogy of RNA regulons with bacterial operons (Keene 2007). In a polycistronic mRNA transcribed from a bacterial operon, the various coding sequences are necessarily found at the same time and place. However, yeast cell wall mRNAs are an example of eukaryotic mRNAs with similar RNA-binding protein complements that are found at different times and places, perhaps more analogous to mRNAs from distinct yet co-regulated and functionally related bacterial operons.

**Figure 5:**
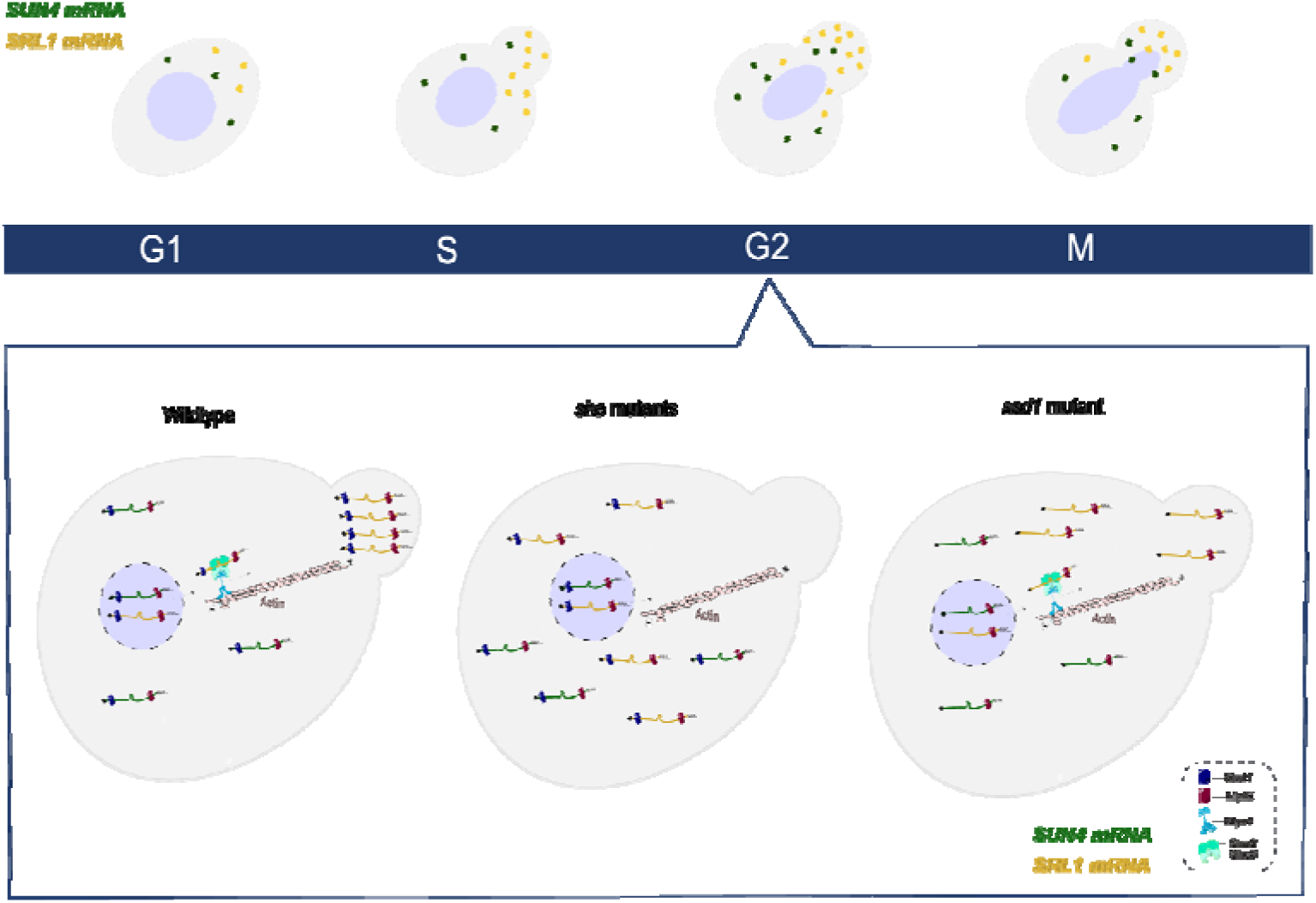
The cell-cycle dependent localization of *SRL1* mRNA to the bud relies on the She2-She3 complex and partially on Ssd1. *SRL1* mRNA is bound by the RBPs Ssd1, She2, and Mpt5. In wild-type, She2 binds *SRL1* mRNA and connects it to the She3-Myo4 protein complex to be shuttled to the budding daughter along actin filaments. In *she* loss-of-function mutants, e.g. *she2*Δ or she3Δ, there is no transportation and localization of *SRL1* mRNA to the bud. Ssd1 deletion partly affects *SRL1* mRNA localization. *SUN4* mRNA is bound by Ssd1 and Mpt5. Mpt5 does not affect localization of either *SUN4* or *SRL1*. Other RNA-binding proteins, although present, are not shown in the figure.

Our observations are consistent with bulk analyses showing that different cwmRNAs have different patterns of expression across the cell cycle (Spellman et al. 1998; Kelliher et al. 2016). The peaks of *SRL1* expression in S/G2 phase and *SUN4* in G1 seen here in smFISH data are also observed in microarray and RNA-seq datasets (Spellman et al. 1998; Kelliher et al. 2016). This reflects the needs of yeast for distinct sets of cell wall proteins at distinct points of the cell cycle, as distinct proteins promote cell wall biogenesis and remodeling through growth of the mother cell, bud emergence and growth, cell septum formation, and then cell separation (Howell and Lew 2012; Orlean 2012). For example, “daughter-specific expressed” mRNAs such as *DSE2* and *CTS1* encode enzymes that remodel the septum from the daughter side (Colman-Lerner et al. 2001). *SUN4* mRNA is most abundant in small, newly separated daughter cells, consistent with its G1 expression peak and with Sun4 acting in cell wall remodeling from the daughter’s side of the septum and in the birth scar (Kuznetsov et al. 2016).

These distinct molecular processes are co-ordinated with both spatial precision on the scale of micrometres or below, and temporal precision on the timescale of minutes (Weiss 2012). Thus, the shared set of RNA-binding proteins that regulates cell walls must support diverse spatiotemporal regulatory programs, rather than enforce spatiotemporal uniformity.

Our results support the conclusion that the She complex confers strong bud localization to mRNAs, that Ssd1 partially contributes to *SRL1* localization, and Mpt5 and Sec72 do not (Figure 6 and Figure S7). For an RBP to be a *bona fide* mRNA localization factor, the simple expectation is that all bound mRNAs are localized and that removing the RBP should abolish mRNA localization. The She complex meets these criteria. Our observation that *SRL1* mRNA is strongly bud-localized, dependent on the She complex, confirms previous results on *SRL1* mRNA and is consistent with the localization of other She-complex bound mRNAs (Shepard et al. 2003). She-dependent binding and localization has been tracked to individual bound “zipcode” fragments within bound mRNAs (Böhl et al. 2000; Cosma 2004; Edelmann et al. 2017; Das et al. 2021), including *SRL1* (Jambhekar et al. 2005). Multiple independent studies have shown that *SRL1* mRNA is bound by She2 (Shepard et al. 2003; Oeffinger et al. 2007; D. J. Hogan et al. 2008; Jambhekar et al. 2005), however evidence that *SUN4* mRNA is bound by She2 comes from only one study (Oeffinger et al. 2007) and no zipcode element has been reported. Overall, the evidence suggests that *SUN4* is not regulated by the She complex.

Our results argue that neither Ssd1 nor Mpt5 are *bona fide* mRNA localization factors, as both these proteins bind some localized mRNAs such as *SRL1*, but also bind some non-localized mRNAs such as *SUN4* (D. J. Hogan et al. 2008; Lapointe et al. 2017; Bayne et al. 2021). The partial role of Ssd1 in localizing *SRL1* mRNA suggests that this protein is not required for active mRNA transport to the bud. Previously, Kurischko and colleagues observed that *SRL1* mRNA bud-localization depended on Ssd1 by using a perturbative labelling method where *SRL1* mRNA was labelled with the MS2 stem-loop and visualized with MS2 coat protein-GFP in live yeast (Kurischko et al. 2011). While we observed a partial disruption of *SRL1* mRNA localization in *ssd1*Δ cells by using smFISH to quantify untagged *SRL1* mRNA in fixed yeast cells, this is qualitatively different from the complete disruption of *SRL1* mRNA localization in *she2*Δ and *she3*Δ strains.

Ssd1 is also associated with other bud-localized mRNAs in yeast including *CLB2* and *MMR1* (Bayne et al. 2021; Hose et al. 2020). Quantification of endogenous mRNA by smFISH shows that deletion of Ssd1 does not affect the localization of pooled *ASH1* + *CLB2* mRNA (Figure 5) or of *CLB2* mRNA alone (Maekiniemi et al. 2025), but does lead to a partial loss of localization of *MMR1* mRNA (Hose et al. 2020). Ssd1 also binds mRNAs such as *DSE2* and *CTS1* (D. J. Hogan et al. 2008; Jansen et al. 2009; Bayne et al. 2021), whose daughter-specific localization is conferred by a distinct transcription-dependent mechanism involving the Ace2 transcription factor (Colman-Lerner et al. 2001). While Ssd1 homologs in filamentous fungi move along hyphal filaments in mRNA-protein complexes (Stein et al. 2020; Modaffari et al. 2026), overall the evidence is weak that Ssd1 binding to mRNAs causes mRNA localization. However, other work suggests a role for Ssd1 in regulation of protein abundance, perhaps via translational regulation (Jansen et al. 2009; Wanless et al. 2014; Kleinerman et al. 2026). An attractive hypothesis to unite these observations would be that Ssd1 works to support She-complex-dependent mRNA transport: while the She complex tethers an mRNP to myosin motors, Ssd1 binding to mRNA could repress translation and so prevent translating ribosomes from removing the She complex from mRNA.

*ASH1* serves as an example where stepwise assembly and remodeling of an mRNP complex enables a specific program that transports an mRNA while repressing translation, followed by localized translational activation (Cosma 2004; Niessing et al. 2018). Fungal cell wall mRNAs, however, must be regulated by a more diverse set of mechanisms than *ASH1*, due to their diverse spatiotemporal localization and diverse set of bound proteins. Therefore, future work needs to mechanistically dissect the regulation of fungal cell wall mRNAs by a system of interacting RBPs, building on quantitative experimental and computational workflows to quantify mRNA localization as presented in this study.

## Materials and Methods

### Yeast strains

BY4741 (MATa his3Δ1 leu2Δ0 met15Δ0 ura3Δ0) was used as wild-type strain (Brachmann et al. 1998). Deletion strains were taken from the yeast genome-wide deletion collection (Giaever et al. 2002) in the BY4741 background with the exception of *she2*Δ, *she3*Δ *and ssd1*Δ, which were generated previously in (Maekiniemi et al. 2025). Yeast strains are described in Table S1. Deletion mutants were verified by PCR.

### Probe design for smFISH

smFISH probes were designed against the native coding sequence of *SUN4* and *SRL1* mRNA using the StellarisTM Probe Designer by LGC Biosearch Technologies, and purchased as pre-Dye labelled probes from Biosearch Technologies. Probe sets were comprised of a set of 39 probes for *SUN4* mRNA, and 25 probes for *SRL1* mRNA (Table S2). Each set of probes were BLASTed against *S. cerevisiae* using the NCBI alignment search tool (https://blast.ncbi.nlm.nih.gov/Blast.cgi). Probes with 17 or more nucleotides complementarity to non-target RNAs were eliminated from the set to prevent off-target binding. Probe sets were labelled with: CAL Fluor Red 610 (CFR610), imaged using the CY3.5 filter for *SRL1* mRNA, and Quasar® 570 (Q570), imaged using the CY3 filter for *SUN4* mRNA. *ASH1* probes and *CLB2* probes, both labelled with Quasar® 670 (Q670), and imaged using CY5 filter, were previously described (Maekiniemi et al. 2025; Tutucci et al. 2018).

### Sample preparation and imaging for smFISH

For smFISH, fixation and hybridization was done following the Tutucci lab protocol (Tutucci and Singer 2020), summarized below. All experiments were conducted in between 3 and 5 biological replicates, i.e. agar plates were streaked with yeast from glycerol stocks, and independent colonies from these plates grown in separate tubes and processed in parallel. All strains were assayed with replicates on multiple different days with consistent results.

### Cell fixation

*S. cerevisiae* strains of interest were grown in Synthetic complete medium (SC medium) + 2% glucose in a 30 °C shaking incubator overnight, then diluted in 25 mL of fresh media and allowed to grow until OD_600_ 0.3–0.4. At this time, cells were fixed through incubation, at room temperature for 45 minutes, with Paraformaldehyde (Thermo Fisher Scientific) at a final concentration of 4% (w/v). Fixed cells were then washed twice with 10 mL of buffer B (1.2 M Sorbitol, 100 mM potassium phosphate buffer [pH 7.5]) at 2400 x g for 3 min at 4°C then resuspended in 1 mL of the same buffer. Cells were digested and permeabilized using lyticase (Sigma-Aldrich) at 25 U lyticase per OD of cells, 20 mM ribonucleoside–vanadyl complex (VRC) (New England Biolabs), and 20mM β-mercaptoethanol and incubated in a water bath at 30 °C for 5-7 minutes. Samples from the incubated cells were checked on a microscope every 2 minutes for cell wall digestion, and removed from water bath when 50% of the cells in the sample are opaque. Following digestion, cells were collected at 1300 x g at 4 °C for 4 min then washed once with 1 mL of buffer B, and resuspended in the same buffer.

### Coverslip preparation

Prior to cell fixation, round noncoated coverslips 0.17 mm thick; diameter 16 mm (Fisherbrand Cover Glasses) were washed with boiling 0.1 M HCl, then autoclaved. Sterile coverslips were stored in 70% Ethanol at 4°C until needed. Coverslips were coated with 0.01% (w/v) poly-L-lysine on the day of the experiment, then placed face-up in sterile 12-well plates using tweezers.

Onto the coated coverslips, ∼125 µL of cells in Buffer B were pipetted to coat each coverslip then incubated at 4 °C for 30 min to an hour. Following the incubation, coverslips were gently washed once with 2 mL of cold Buffer B, with the buffer carefully removed using a vacuum aspirator. 2 mL of cold 70% ethanol was gently added to each well for storage. Plates were sealed with parafilm and stored at-20 °C overnight or until ready to be used for probe hybridization.

### Probe hybridization

For hybridization, coverslips were placed in a new 12-well plate and rehydrated with two washes of 2 mL 2x Saline sodium citrate (SSC) (diluted from 20x SSC: 3 M NaCl, 0.3 M sodium citrate-HCl, pH 7.0). Following these washes, coverslips were incubated for 30 min in 2 mL of 10% formamide/2xSSC at room temperature. For each coverslip: 0.125 µL of the probe stock (stock concentration is 25 µM) is needed to achieve a final concentration of 125 nM in 25 µL of the hybridization mix. The probe mix was prepared for 4 coverslips by taking 0.5 µL of each probe and adding to 20 µL (5 µL per coverslip) of *E. coli* tRNA/ssDNA (10 mg/mL) in a 1.5 mL Eppendorf tube. The probe mix was lyophilized in a SpeedVac at 45 °C, to be resuspended in the hybridization mix later, and protected from light by using foil to cover the tubes.

Following the lyophilization step, the hybridization mix was added to the probe mix in two steps. First, a solution containing 20% formamide and 10 mM NaHPO4 [pH 7.5] was added to the probe mix tube, at 12.5 µL per coverslip, then incubated at 95 °C for 2 min then cooled at RT in the dark. Following this, 12.5 µL per coverslip of a solution containing 4xSSC, 2 mg/mL BSA, and 10 mM VRC of was added to the same tube. For each coverslip, 20 µL of the hybridization mix was pipetted onto parafilm placed in a hybridization chamber (square plate covered in foil). Using tweezers, coverslips were carefully placed on the hybridization mix with the cells facing down, then incubated at 37 °C for 3 hours. A plastic cap filled with water was placed inside each hybridization chamber to create humidity.

Coverslips were placed back in the 12-well plates and washed twice with 2 mL of 10% formamide/2xSSC and incubated at 37 °C for 15 min each time. Following this, coverslips were washed once with 2 mL 2xSSC 0.1% Triton X-100 at RT (5 min), once with 2 mL of 2xSSC at RT (5 min), and a final wash with 2 mL of 1xSSC at RT (5 min). During the last incubation, glass slides were labelled with date, initials, and the strain name plus probe set used, then placed inside clean hybridization chambers. Coverslips were washed by dipping them in 100% ethanol and let them airdry while protecting them from ambient light. One drop of ProLong Diamond Antifade Mountant with DAPI (Thermo Fisher Scientific) was placed on each glass slide. Using tweezers, coverslips were transferred from the plate and onto the glass slide with cells facing down. Slides were incubated at RT overnight in the dark in the hybridization chambers and sealed with clear nail polish in the morning. Following this, slides can be stored in the - 20 °C for months.

### Combined smFISH and immunofluorescence (smFISH-IF)

The smFISH-IF images in Figure 1B were acquired in a previous study and first reported in a preprint (Maekiniemi et al. 2022). Briefly, after the final smFISH wash, samples were post-fixed in 4% PFA in 1× PBS for 10 min at room temperature (RT) and washed for 5 min in 1× PBS. Coverslips were blocked for 30 min at RT in 1× PBS containing 0.1% RNase-free bovine serum albumin (BSA) and then incubated for 45 min with a mouse anti-tubulin antibody (Thermo Fisher, 1:1000) in the same buffer. After a 5-min wash in 1× PBS, coverslips were incubated for 45 min at RT with goat anti-mouse Alexa Fluor 488 (1:1500) in 1× PBS, 0.1% BSA, and washed three times for 5 min in 1× PBS. Coverslips were dehydrated in 100% ethanol, air-dried and mounted with ProLong Gold Antifade Mountant with DAPI (ThermoFisher).

### Imaging of fixed cells

For smFISH imaging, we used an Olympus BX63 epifluorescence microscope equipped with an Ultrasonic stage and UPlanApo 100X 1.35NA oil-immersion objective (Olympus). CellSens (Olympus) software was used for instrument control and image acquisition. Lumencore SOLA FISH light source, a Hamamatsu ORCA-Fusion sCMOS camera (pixel-size 6.5 µm x 6.5 µm) mounted using U-CMT C-Mount Adapter, and zero-pixel shift filter sets: F36-500 DAPI HC Brightline Bandpass Filter, Filter, F36-542 Cy3 HC BrightLine Filter (for Q570-labelled probes), AHF-LED-FISH-R (for CFR610-labelled probes), and F36-523 Cy5 HC BrightLine Filter (for Q670-labelled probes), and a single plane differential interference contrast (DIC) image for 50–100 ms. For the DAPI channel 25–50 ms exposure was used for each Z plane, while for smFISH channels an exposure time of 750 ms was used to acquire each Z plane. Images are acquired across 41 optical sections (or adjusted, depending on the sample thickness) with a Z-step size of 0.2 μm.

### Image analysis

RNA quantification and assignment of mother–bud relations were performed using custom Python packages, buddy and napari-buddy. These packages act as a wrapper around the Big-FISH spot analysis package (Imbert et al. 2021), the Omnipose cell-segmentation package (Cutler et al. 2022), and the Python image viewer napari (napari contributors 2019). The analysis takes the Olympus VSI format as input. In brief, it follows these steps: cell segmentation from the DAPI and DIC layers with Omnipose; spot detection and decomposition with Big-FISH; data visualization and mask correction in napari; manual mother–bud assignment in napari aided by napari-buddy; and cell-cycle assignment based on the extracted features (also see Figure 1).

In more detail, cell masks were segmented with the Omnipose cyto2_omni model using the custom parameters: channels = [1, 2], mask_threshold = −3, flow_threshold = 1, min_size = 10, diameter = 50, and cluster = True; all other parameters were kept at their defaults. The model was applied to an overlay of the max-projected DAPI channel and the DIC image. Nuclear masks were segmented with the nuclei model on the max-projected DAPI channel, using the parameters: channels = [0, 0], mask_threshold = 0, flow_threshold = 0, min_size = 10, diameter = 0, and cluster = True. Cell masks without a nucleus and masks smaller than 20 pixels were removed, as were nuclear masks lying outside a cell.

To determine the RNA-quantification parameters for a batch of images from a single FISH experiment, a representative image was first loaded into napari-buddy, and its background-filtering and spot-detection UI elements were used to set the detection parameters. Background was removed with a Gaussian filter (Big-FISH remove_background_gaussian; sigma [z:0.75, y:2.3, x:2.3]), after which spots were detected with the Big-FISH detect_spots function, which applies a Laplacian-of-Gaussian filter followed by local-maximum detection and thresholding. Voxel size (z:200, y:65, x:65 nm), spot radius (z:880, y:120, x:120 nm) and sigma were kept the same across the entire dataset, whilst the spot-detection threshold was adapted per experiment to account for differences in signal intensity. Dense spot regions were decomposed with the Big-FISH decompose_dense function (α = 0.5, β = 2, γ = 1) to estimate the number of RNAs they contained. These parameters were saved to a config.json file via the ‘Create config file’ UI element and reused to batch-process all images of that experiment.

Batch segmentation, spot detection and spot decomposition were performed in the buddy Jupyter notebook ‘01_fish_bulk_processing.ipynb’. To speed up spot detection, the in-focus z-range was determined for each image and fluorescence channel on a [200 × 200]-pixel patch with the highest cell density. The Big-FISH focus measure was computed per slice, max-projected and smoothed with a Savitzky–Golay filter (window of 16 slices, second-order polynomial). The in-focus range was defined as the slices between the steepest rise and steepest fall of this profile, extended by 5 slices below and 10 slices above (median 27 slices, range 16–39; ∼5.4 µm, range 3.2–7.8 µm). Spots were detected only within this range, and spots in its lowest three slices were discarded, as these contain probes attached directly to the coverslip glass. Spot decomposition was performed subsequently with parameters described above.

The DIC and fluorescence channels were offset by ∼5–15 pixels, owing to a shift introduced by the DIC filter in the light path. To correct this, spots from all mRNA channels were pooled, and the translation that best placed them within cells was estimated by a custom algorithm. For each pixel, the distance to the nearest cell mask was computed (capped at 50 pixels), and each candidate shift was scored by the mean distance of the shifted spots. A coarse Monte Carlo search over 4,000 random integer shifts (up to ±200 pixels in x and y) was followed by an exhaustive local search within ±20 pixels of the best candidate. The inverse of the resulting translation was applied to the DIC image and cell masks using nearest-neighbour interpolation. Spot coordinates and nuclear masks, both in the fluorescence frame, were left unchanged, and the applied shift was recorded in each image’s metadata. For part of the data, masks were aligned manually, because this tool was developed while the data were being processed.

The resulting spot detections and aligned masks were then inspected in napari, where cell and nuclear masks could be manually corrected if necessary. A mask describing mother–bud relations was then created manually using the napari-buddy ‘Reassign buds to mothers’ UI element, which lets users draw lines linking mothers and buds.

After the manual curation was complete, two bud-detection models were trained on the curated bud masks (buddy ≥ 0.4.0, bud_model_DIC_DAPI_only and bud_model_with_ASH1CLB2; notebook 05_train_bud_model.ipynb). The 132 imaged fields were split whole into training (66 fields), validation (6) and test (20) sets so that no cell of a test field was seen during training; the 21 fields of imaging day 220429 (poor DIC quality resulting in poor Omnipose segmentation) and the remaining 19 fields of the test day were not used (she-mutant strains or curation was not checked for missing small buds). Each cell segmented by Omnipose is presented to the model as a 192 × 192 px (12.5 µm) crop centred on the cell, with the channels DIC, maximum-projected DAPI and a drawn dot marking the queried cell, plus the maximum-projected CLB2 + ASH1 channel for the second model (trained without the she2Δ and she3Δ strains, in which this channel is uninformative). The network is a U-Net with a ResNet-34 encoder pretrained on ImageNet (segmentation_models_pytorch; 24.4 M parameters) with two output maps (bud of this cell, body of this cell; binary cross-entropy + Dice loss) and a three-class head on the encoder (unbudded, budded, bud; cross-entropy). Models were trained for 30 epochs (batch 16), and the last epoch was used. A cell’s bud is accepted when P(budded) × mean bud-map probability ≥ 0.5; when several cells claim the same bud the strongest claim wins. Performance (Figure S2) was measured on the 14 test fields of imaging day 240909 (wild type, ssd1Δ, sec72Δ; 2,361 cells, 348 buds) as mother–bud pair F1 (a predicted bud counts as correct when it overlaps the annotated bud of the same mother), with 95% confidence intervals from 1,000 bootstrap resamples over cells; 43 cells whose annotation was judged uncertain on review were excluded.

Spots and decomposed dense regions were assigned to cells, buds and nuclei based on their x–y position in the corresponding masks. The total number of RNAs per cell was calculated as the number of single spots plus the estimated number of RNAs in dense regions. Dense regions overlapping a nucleus were classified as transcription sites, and the RNAs they contained as nascent RNAs. Counts in the mother compartment were obtained by subtracting bud counts from whole-cell counts. Nuclei were assigned to the mother or bud by the position of their centroid, and per-compartment nuclear counts, together with cell and bud morphology (area, axis lengths, eccentricity, solidity), were exported for each cell to a.csv file.

Lastly, cell-cycle phases were assigned automatically based on bud area, the number of nuclei per compartment, and the number of *ASH1* + *CLB2* transcripts. Pixel measurements were converted to micrometres (0.065 µm per pixel), and only cells with a major-axis length between 2 and 20 µm were considered. All cells required a single nucleus in the mother, and were classified as follows: G1, no bud and five or fewer *ASH1* + *CLB2* transcripts; S, bud area below 2 µm² with no nucleus in the bud; G2, bud area of at least 2 and below 15 µm² with no nucleus in the bud; and M, bud area above 2 µm² with a single nucleus in the bud. Cells meeting none of these criteria, such as unbudded cells with more than five *ASH1* + *CLB2* transcripts, cells with buds of 15 µm² or larger lacking a bud nucleus, or cells without exactly one mother nucleus, were left unassigned.

To validate the automatic cell-cycle assignment, a subset of assigned cells from all strains were manually re-annotated. Each cell was exported as a cell-centred multi-channel TIFF containing the fluorescence channels, the max-projected DAPI channel, the aligned DIC image, and a dashed outline marking the cell of interest, with fixed per-channel display ranges; crops were organized by their automatically assigned phase. Up to 100 crops were generated per phase (390 in total: 99 G1, 96 S, 97 G2, 98 M). Four validators each independently scored a random sample of 100 crops while blinded to the automatic call, covering 271 unique cells (400 annotations in total, with partial overlap between annotators). Cells marked “Unsure” (n = 28) were excluded, leaving 372 annotations. Agreement between the automatic assignment and the manual annotation was quantified as the overall percentage of matching calls and Cohen’s κ, with per-phase performance summarised as precision (fraction of automatic calls confirmed manually) and recall (fraction of manually annotated cells recovered). The automatic assignment agreed with manual annotation for 84.7% of annotations (Cohen’s κ = 0.80), with consistent agreement across annotators (82–87%). Agreement was highest for G1 and M (precision 94% and 90%) and lowest for S (precision 72%), which was most often confused with the adjacent G1 and G2 phases.

Linking buds to mothers was the main manual effort in this workflow (2,406 buds). After completing the analysis, we therefore used the curated dataset to train a neural network that performs this step automatically (Figure S2; Methods). Model performance was estimated on unseen test data, acquired on a separate day as any of the training data (Figure S2; Methods). For every segmented cell, a U-Net (Ronneberger et al. 2015) with a ResNet-34 encoder receives a 192 × 192 px crop of the DIC and DAPI channels, with a dot marking the cell in question, and returns the bud of that cell, when present. A second trained model additionally receives the CLB2 + ASH1 channel during training. The DIC + DAPI model recovered 72% of the annotated buds with a precision of 80%, the CLB2 + ASH1 model reached a precision of 91% at the same recall. The models are available with buddy as an optional step that automatically segments buds for the user to check in napari-buddy, further speeding up the analysis.

Validation analyses were performed in Python using pandas (McKinney 2010), scikit-learn (Pedregosa et al. 2012) and matplotlib (Hunter 2007), and all other data were analyzed and plotted in R (R Core Team 2023) using tidyverse (Wickham et al. 2019), ggplot2 (Wickham 2009), cowplot (Wilke 2025), and ggsignif (Ahlmann-Eltze and Patil 2021).

## Data availability

The image processing pipelines buddy and napari-buddy are available on Zenodo doi: 10.5281/zenodo.23771337

The data and code underlying quantitative analysis in Figures 2, 3, 4, S3, S4, S5, S6, and S7 is available on GitHub at https://github.com/ewallace/cwmrnaloc_2026 and Zenodo at doi:10.5281/zenodo.24315446.

## Author Contributions

Conceptualization: E.W.J.W., E.T., and M. M. J. Data curation: E.W.J.W., M. M. J., and S. v. O. Formal analysis: E.W.J.W., C.M., M. M. J., S. v. O., and K. v. R. Investigation: M. M. J., K. v. R., and S. v. O. Methodology: E.W.J.W., E.T., M. M. J., S. v. O., C. M., and P.S. Software: S. v. O., C.M., F.B., and P.S. Supervision: E.W.J.W. and E.T. Validation: S. v. O., K. v. R., and E.T. Visualization: E.W.J.W., E.T., M. M. J., and S. v. O. Writing—original draft: E.W.J.W., E.T., M. M. J., and S. v. O. Writing—review and editing: all.

## Acknowledgments

We thank Atlanta Cook, Liz Ballou, Amy Buck, Stefan Bresson, David Tollervey, Ivan Clark, and members of the Wallace and Tutucci groups for extensive discussions relating to the project. We thank Jean Beggs and Edinburgh Genome Foundry for providing strains. We thank Sabine Michielsen, Luna Meister and Dennis Botman (Vrije Universiteit Amsterdam) for help with validation experiments.

