## Supplementary Material (Supplementary Figures S1-S5, Supplementary Tables S1-S2). for "Multi-colour smFISH with automated cell-cycle quantification reveals distinct localization of co-regulated cell wall mRNAs in *Saccharomyces cerevisiae* yeast"

**Contents:**

**Supplementary Figures S1-S5**

**Figure S1: Validation of automatic cell-cycle assignment against manual annotation.**

**Figure S2: Automatic bud detection by U-net-based segmentation.**

**Figure S3: Images of representative cells from phase G1 in wild-type (BY4741), *ssd1∆*, *mpt5∆*, *she2∆* and *she3∆* cells.**

**Figure S4: *SRL1* mRNA localization in M phase is She2- and She3-dependent.**

**Figure S5: Statistical analysis of mRNA bud-localization of *CLB2 + ASH1*, *SRL1*, and *SUN4* mRNA.**

**Figure S6:** **Nascent mRNA transcription in the *SRL1* and *SUN4* mother and bud nuclei is tightly coordinated with cell size.**

**Figure S7: *SRL1* and *SUN4* mRNA localization is not dependent on the Sec complex*.***

**Supplementary Tables**

**Table S1. Yeast Strains used in this study.**

**Table S2. smFISH probe sequences.**

**Supplementary figures**

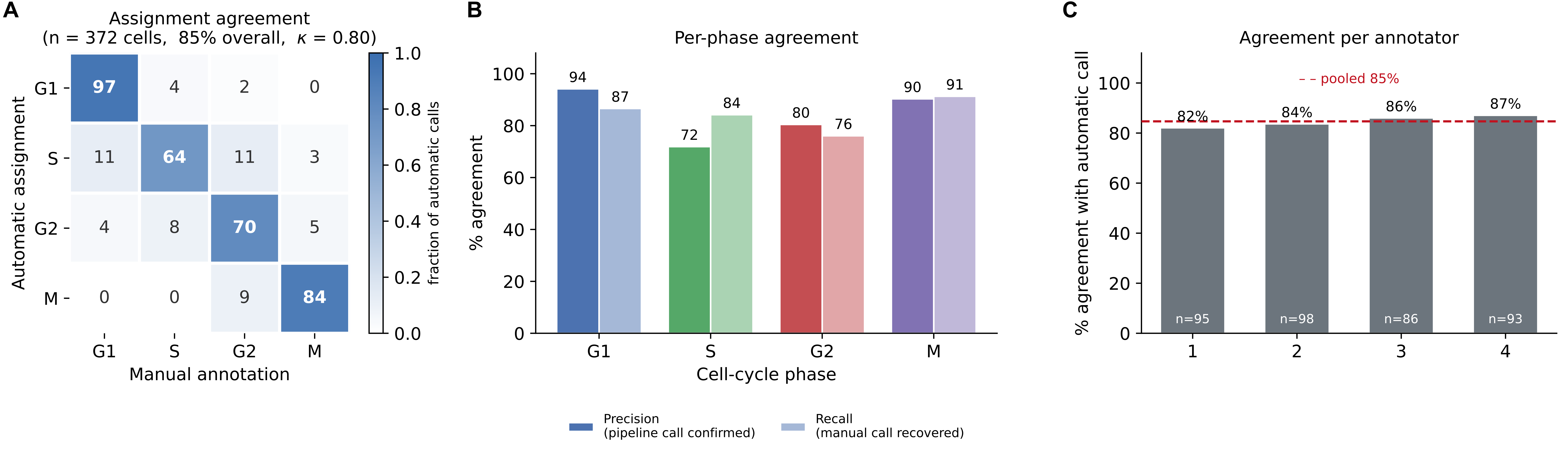

**Figure S1: Validation of automatic cell-cycle assignment against manual annotation.** Related to Figure 1. Four annotators independently re-assigned crops from a pool of 390 automatically assigned cells (~100 per phase) randomly chosen from our entire datasets, yielding 372 annotations after exclusion of "Unsure" calls. (A) Confusion matrix comparing automatic assignment (rows) with manual annotation (columns); cells are coloured by the fraction of automatic calls per row, with counts overlaid (overall agreement 85%, Cohen's κ = 0.80), where κ = (*p*ₒ − *p*ₑ) / (1 − *p*ₑ). Here *p*ₒ is the observed agreement and *p*ₑ is the agreement expected by chance, calculated from how often each method assigns each phase (*p*ₑ = Σₖ *p*ₖ,auto · *p*ₖ,manual). κ = 0.80 indicates that the automatic assignment captures 80% of the agreement with manual annotation that is not attributable to chance. (B) Per-phase precision (fraction of automatic calls confirmed manually) and recall (fraction of manually annotated cells recovered). (C) Agreement with the automatic call per annotator; the dashed line marks the pooled value, showing consistency across validators.

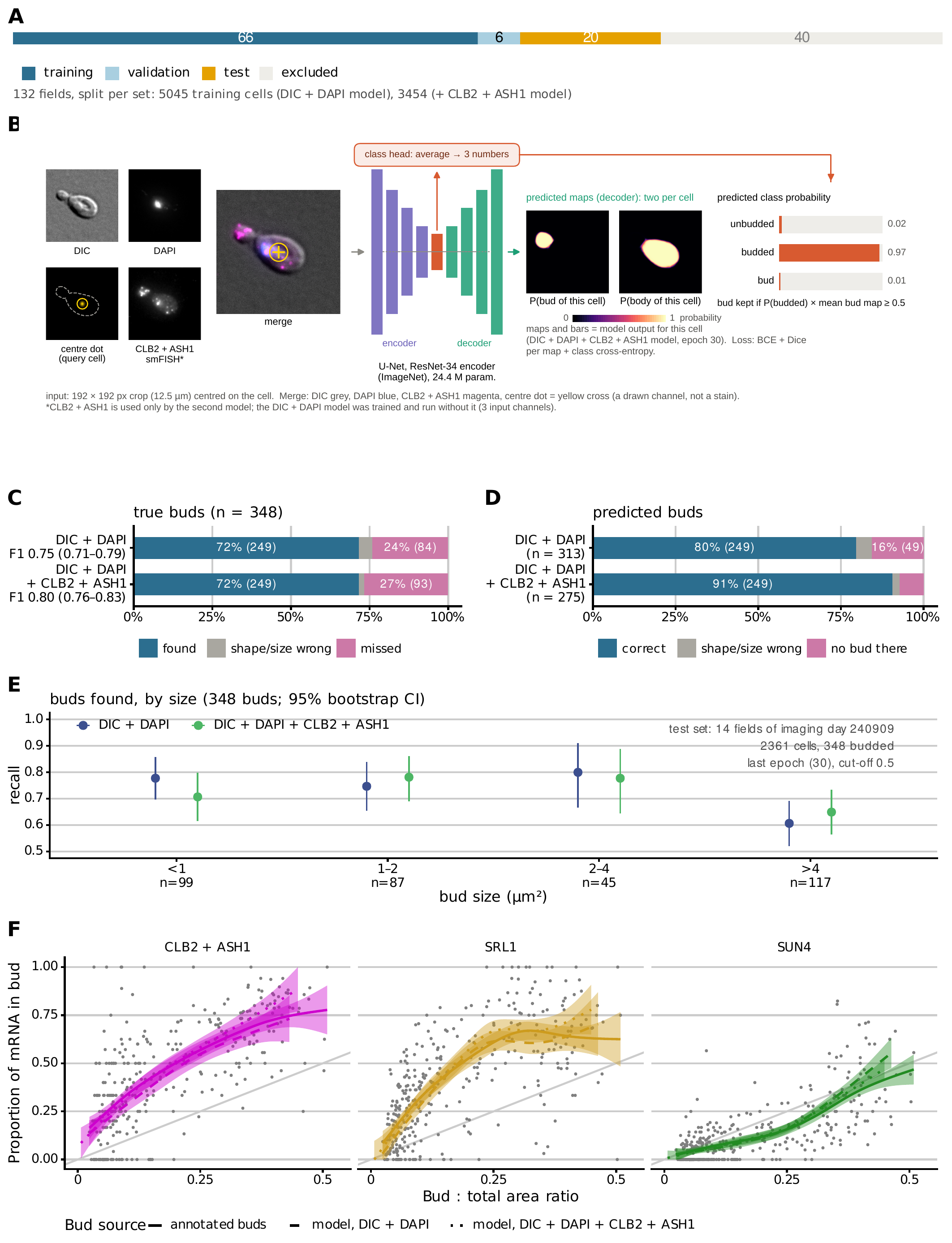

**Figure S2: Automatic bud detection by U-net-based segmentation.** **(A)** Data split. The 132 imaged fields (6 imaging days, 7 strains) were assigned whole to the training (66), validation (6) and test (20) sets, so that no cell of a test field was ever seen in training; 40 fields were excluded (the other 19 fields of the test day 240909 and the 21 fields of day 220429 with poor DIC). The training fields hold 5045 annotated cells for the DIC + DAPI model and 3454 for the DIC + DAPI + CLB2 + ASH1 model, which was trained without the *she2*∆ and *she3*∆ strains. **(B)** The per-cell model. Every segmented cell is queried once with a 192 × 192 px crop (12.5 µm) centred on it, with the channels DIC, DAPI, a drawn centre dot marking the queried cell and, for the second model, the CLB2 + ASH1 smFISH channel. A U-Net (ResNet-34 encoder pretrained on ImageNet, 24.4 M parameters) returns two probability maps at crop resolution, the bud of this cell and the body of this cell, and, from a classification head on the encoder, the probabilities of the three classes unbudded, budded and bud. A bud is kept when P(budded) × mean bud-map probability ≥ 0.5; when several cells claim the same bud the strongest claim wins. Shown: a real cell of the test day (wild type, 3.3 µm² bud) with the output of the DIC + DAPI + CLB2 + ASH1 model, P = 0.02 / 0.97 / 0.01. **(C–E)** Bud detection on an unseen imaging day: the 14 test fields of day 240909 (wild type, *ssd1*∆, *sec72*∆; 2361 cells, 348 budded), every cell annotated by hand, both models at their last epoch (30) and cut-off 0.5, fixed before scoring. **(C)** The 348 true buds: found with the right shape and size (overlap with the annotation above the cut-off), found with the wrong shape or size, or missed, for each model; pair F1 with its 95 % bootstrap confidence interval over cells. **(D)** The buds predicted by each model: correct, wrong shape or size, or on a cell without a bud. **(E)** Recall of the true buds by bud size, 95 % bootstrap confidence intervals; n = buds per size class. **(F)** The biology is the same with model buds. Proportion of mRNA in the bud against the bud : total area ratio for the three mRNAs, on the budded cells of the test fields (all three strains pooled), with the buds taken from the annotation (solid line, grey points) or predicted by the DIC + DAPI model (dashed) or the DIC + DAPI + CLB2 + ASH1 model (dotted). Lines are GAM smooths with their standard-error bands, as in Figure 4B; the grey line y = x is the proportion expected if the mRNA were spread evenly over the cell. Cells with at least one mRNA of that species: annotated buds n = 342, model buds n = 307 and 270.

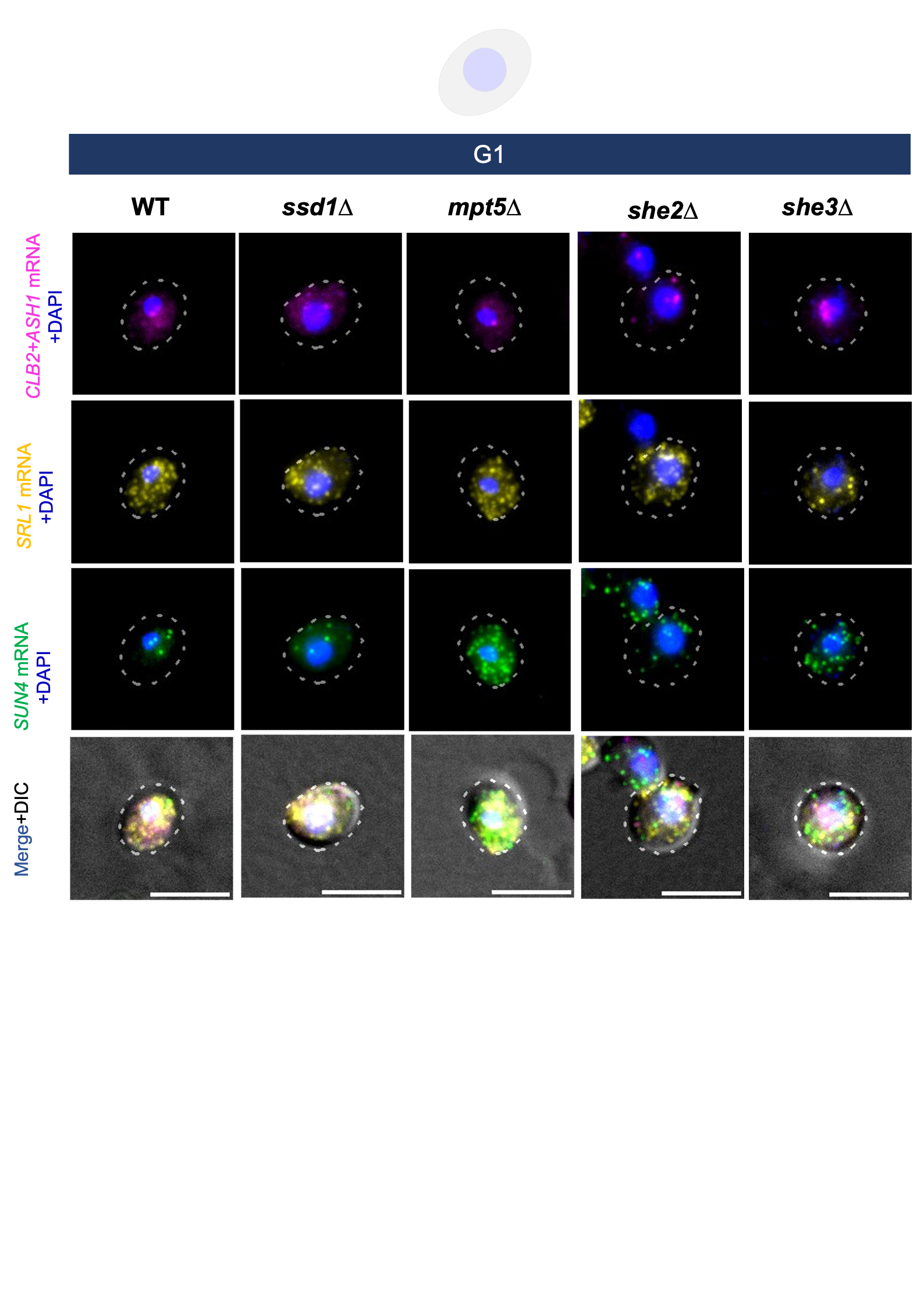
 **Figure S3: Images of representative cells from phase G1 in wild-type (BY4741), *ssd1∆*, *mpt5∆*, *she2∆* and *she3∆* cells.** From top to bottom, smFISH images of *CLB2 + ASH1* mRNA (magenta), *SRL1* mRNA (yellow), *SUN4* mRNA (forest green), and merged images with all three mRNAs and DIC. All images show the cell outline in dashed white line, and the nucleus stained by DAPI in blue, scale bars are 5 μm. Related to Figure 3.

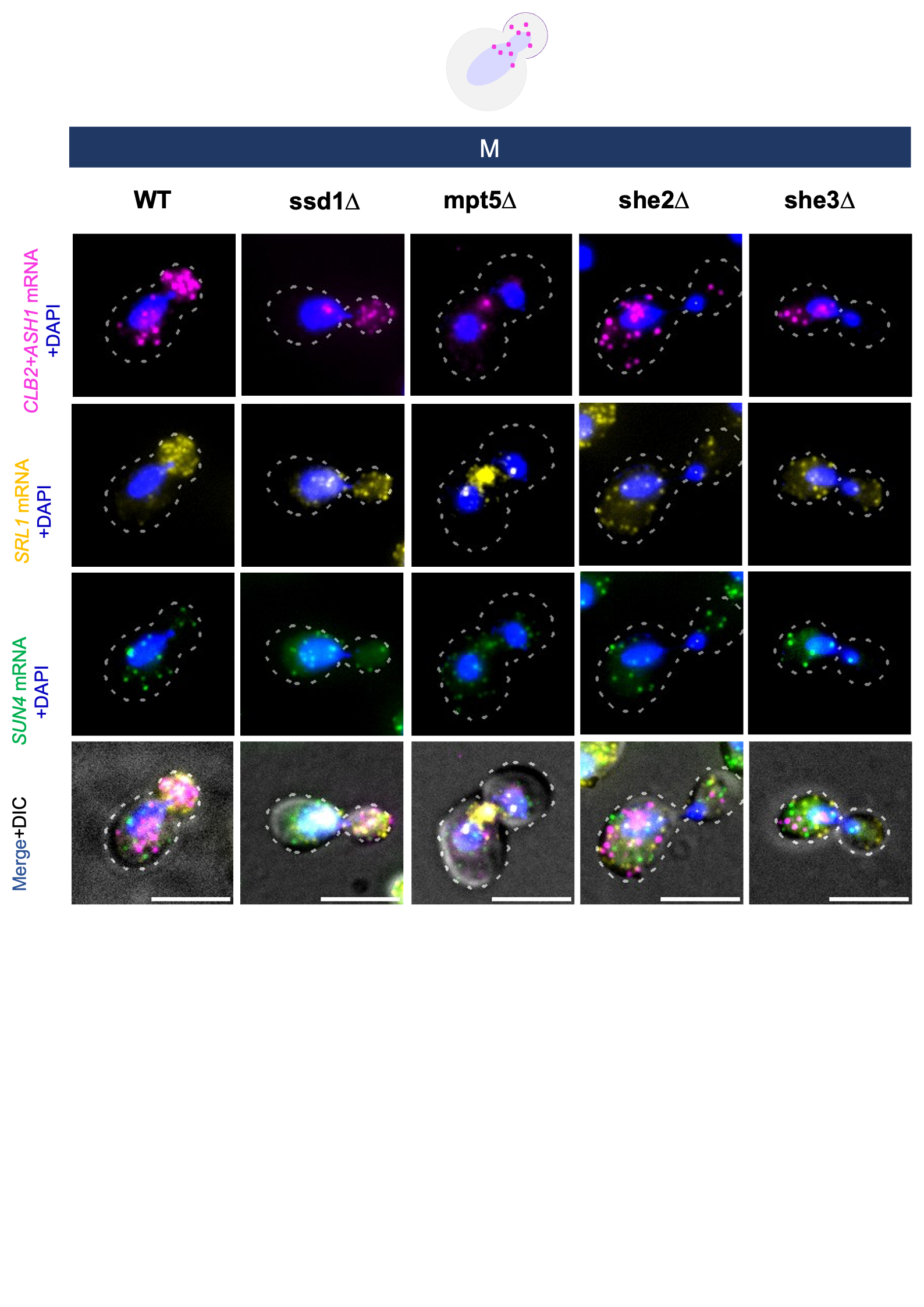

**Figure S4: *SRL1* mRNA localization in M phase is She2- and She3-dependent.** Images of representative cells from phase M in wild-type (BY4741), *ssd1∆*, *mpt5∆*, *she2∆* and *she3∆* cells; from top to bottom, smFISH images of *CLB2 + ASH1* mRNA (magenta), *SRL1* mRNA (yellow), *SUN4* mRNA (forest green), and merged images with all three mRNAs and DIC. All images show the cell outline in dashed white line, and the nucleus stained by DAPI in blue, scale bars are 5 μm. *SRL1* mRNA localizes to the bud in wild-type cells. In comparison, *SRL1* mRNA bud-localization is abolished in *she2*∆ and *she3*∆ cells, similarly to positive-control mRNAs *CLB2 + ASH1,* and partially maintained in *ssd1∆* cells. Related to Figure 3.

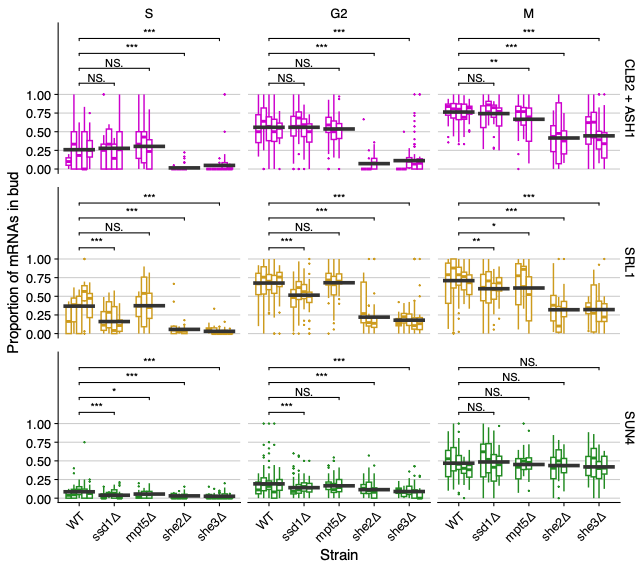

**Figure S5: Statistical analysis of mRNA bud-localization of *CLB2 + ASH1*, *SRL1*, and *SUN4* mRNA.** Related to Figure 3. Each boxplot represents one biological replicate. The black horizontal bars show the mean proportion of RNA in bud across all replicates. The brackets and significance values compare mean proportion in bud between wild-type and deletion strains, calculated using the t-test as implemented in the lm function in the stats package in R (R Core Team 2023), and visualised using the ggsignif package (Ahlmann-Eltze and Patil 2021).

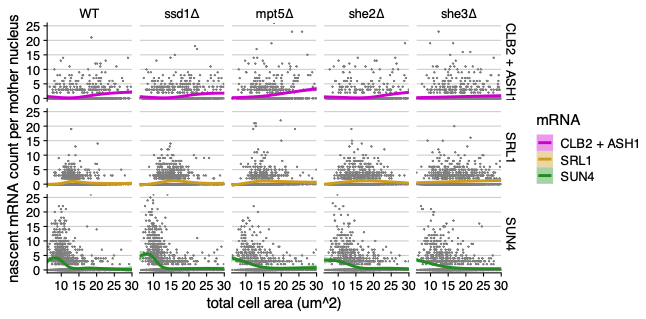

**Figure S6:** **Nascent mRNA transcription in the *SRL1* and *SUN4* mother and bud nuclei is tightly coordinated with cell size.** Related to Figure 4. The y-axis, nascent mRNA count per mother nucleus, is the number of mRNAs estimated at the site of transcription in the mother cell nucleus from smFISH data (see methods).

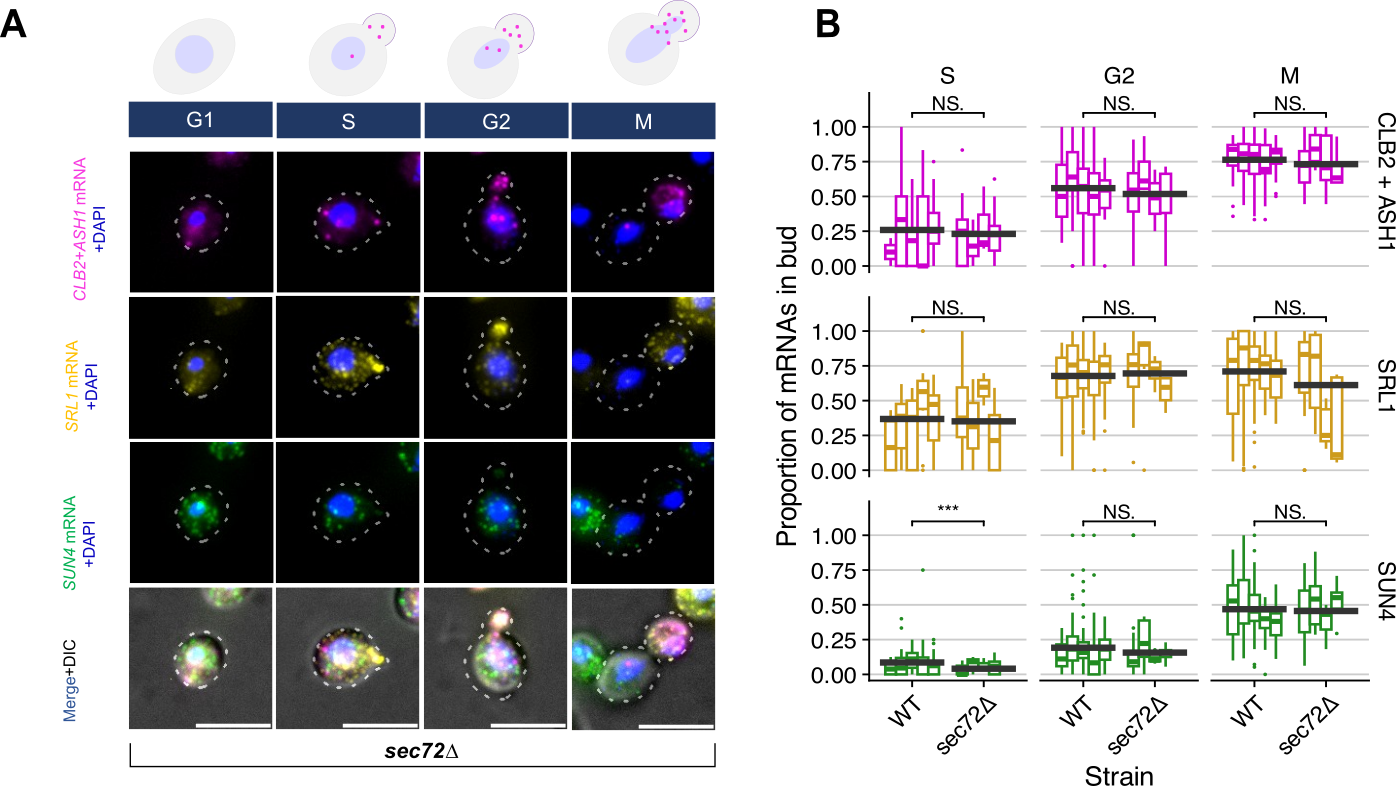

**Figure S7:** ***SRL1* and *SUN4* mRNA localization is not dependent on the complete Sec complex*.*** **A**, Images of cells from phases of the cell cycle (G1, S, G2, and M) in *sec72∆*; from top to bottom, smFISH images of *CLB2 + ASH1* mRNA (magenta), *SRL1* mRNA (yellow), *SUN4* mRNA (forest green), and merged images with all three mRNAs and DIC. All images show the cell outline in dashed white line, and the nucleus stained by DAPI in blue, scale bars are 5 μm. In these images, *SRL1* mRNA and positive control *CLB2 + ASH1* mRNA localization to the bud is unaffected by the Sec complex mutant *sec72∆. SUN4* mRNA is also unaffected. **B**, Quantification of proportion of mRNA in bud across replicates confirms that *SRL1* mRNA localization is unaffected in *sec72∆*. There is a small but statistically significant decrease in the proportion of *SUN4* mRNA in the bud in S-phase cells, in *sec72∆* compared to wild-type.

| **Code** | **Name** | **Description** | **Background** | **Genotype** | **Source** |
| --- | --- | --- | --- | --- | --- |
| YET563 | *she2∆* | ΔYKL130C | BY4741 | *MATa;his3Δ1; leu2Δ0; met15Δ0; ura3Δ0 YKL130C:: HygB* | (Maekiniemi et al. 2025) |
| YET572 | *she3∆* | ΔYBR130C | BY4741 | *MATa;his3Δ1; leu2Δ0; met15Δ0; ura3Δ0 YBR130C:: KANr* | (Maekiniemi et al. 2025) |
| YET915 | BY4741 | Wild-type | S288C | *MATa; his3Δ1; leu2Δ0; met15Δ0; ura3Δ0;* | (Brachmann et al. 1998) |
| YET916 | *ssd1∆* | Δ YDR293C | BY4741 | *MATa; his3Δ1; leu2Δ0; met15Δ0; ura3Δ0; YDR293C::kanMX4* | Deletion collection (Giaever et al. 2002). |
| YET919 | *mpt5∆* | ΔYGL178W | BY4741 | *MATa; his3Δ1; leu2Δ0; met15Δ0; ura3Δ0; YGL178W::kanMX4* | Deletion collection (Giaever et al. 2002). |
| YET921 | *sec72∆* | ΔYLR292C | BY4741 | *MATa; his3Δ1; leu2Δ0; met15Δ0; ura3Δ0; YLR292C::kanMX4* | Deletion collection (Giaever et al. 2002). |

**Table S1. Yeast Strains used in this study.** We thank Jean Beggs for assistance obtaining BY4741. We thank Edinburgh Genome Foundry for the yeast deletion collection strains from (Giaever et al. 2002).

| GENE | Probe # | SEQUENCE | Dye |
| --- | --- | --- | --- |
| SRL1 | 1 | atcgctgacacagaacttgc | CAL Fluor Red 610 |
| SRL1 | 2 | gcgctaaagtggtagttgta | CAL Fluor Red 610 |
| SRL1 | 3 | cttggggcaccaaggagtag | CAL Fluor Red 610 |
| SRL1 | 4 | gtcggcgtacgatatggtag | CAL Fluor Red 610 |
| SRL1 | 5 | gtagaccgttgaggtgacaa | CAL Fluor Red 610 |
| SRL1 | 6 | ttgaggtgaaccagctcgtg | CAL Fluor Red 610 |
| SRL1 | 7 | gcattggtaatggtggctga | CAL Fluor Red 610 |
| SRL1 | 8 | ggtggacaaggaggaggagg | CAL Fluor Red 610 |
| SRL1 | 9 | ttacagatccagaggccgaa | CAL Fluor Red 610 |
| SRL1 | 10 | atttcatgggtggattctgg | CAL Fluor Red 610 |
| SRL1 | 11 | cgtgatagtcgaggtggagg | CAL Fluor Red 610 |
| SRL1 | 12 | gaagggttagcagcaaagtg | CAL Fluor Red 610 |
| SRL1 | 13 | ggagacaaagtagtggagtc | CAL Fluor Red 610 |
| SRL1 | 14 | gacacttgctgcagtagatg | CAL Fluor Red 610 |
| SRL1 | 15 | tgttgtttgaatcttcgtca | CAL Fluor Red 610 |
| SRL1 | 16 | aaggacttgacctttgcatc | CAL Fluor Red 610 |
| SRL1 | 17 | cattggaagttgaagcctgt | CAL Fluor Red 610 |
| SRL1 | 18 | aactttgtgattgggacgca | CAL Fluor Red 610 |
| SRL1 | 19 | taacgggctcattggtgaca | CAL Fluor Red 610 |
| SRL1 | 20 | ttggggtgactgtaacgtac | CAL Fluor Red 610 |
| SRL1 | 21 | gtaacgtattgtgtagtcgt | CAL Fluor Red 610 |
| SRL1 | 22 | taacagaaggtgcaccggtg | CAL Fluor Red 610 |
| SRL1 | 23 | tacgttacctggagaggtag | CAL Fluor Red 610 |
| SRL1 | 24 | tcgaagtggtgttgtaccat | CAL Fluor Red 610 |
| SRL1 | 25 | caccaactggtcgaattagt | CAL Fluor Red 610 |
| GENE | Probe # | SEQUENCE | Dye |
| SUN4 | 1 | caatgaagccgcggtaagag | Quasar 570 |
| SUN4 | 2 | agcggaaacgattgtgctat | Quasar 570 |
| SUN4 | 3 | tcaatatcagccgcatacgg | Quasar 570 |
| SUN4 | 4 | agcagtagtagtacatcctg | Quasar 570 |
| SUN4 | 5 | tcttatgctggtgtgaacca | Quasar 570 |
| SUN4 | 6 | acgtatgtaactgcaacggc | Quasar 570 |
| SUN4 | 7 | atcgacagtgactgtttcgt | Quasar 570 |
| SUN4 | 8 | gagtgacggtctgaccattt | Quasar 570 |
| SUN4 | 9 | agaagaggcttcagttgagg | Quasar 570 |
| SUN4 | 10 | tcagagtagtagtggaagca | Quasar 570 |
| SUN4 | 11 | acttggtgaccgaagattcg | Quasar 570 |
| SUN4 | 12 | agcgatttgttcagtactct | Quasar 570 |
| SUN4 | 13 | tttgagccgaagacgaggtg | Quasar 570 |
| SUN4 | 14 | cacagaggattcgctagtag | Quasar 570 |
| SUN4 | 15 | ctgacccagatgtggaaata | Quasar 570 |
| SUN4 | 16 | gaagctgctgaagatgtgga | Quasar 570 |
| SUN4 | 17 | atggacctgaaaagtcagcc | Quasar 570 |
| SUN4 | 18 | aaggaatggtaccgtcttca | Quasar 570 |
| SUN4 | 19 | ttcatctaaccagctgatgg | Quasar 570 |
| SUN4 | 20 | gtattttcaacaccagacca | Quasar 570 |
| SUN4 | 21 | tacaagagccacccgtagaa | Quasar 570 |
| SUN4 | 22 | tatgagcagtaagacccttc | Quasar 570 |
| SUN4 | 23 | cattgtgtcttggacatacc | Quasar 570 |
| SUN4 | 24 | gtcagatggttggtcagaag | Quasar 570 |
| SUN4 | 25 | tttgaacggtacaggtagcc | Quasar 570 |
| SUN4 | 26 | cccattcgcataaatagtca | Quasar 570 |
| SUN4 | 27 | ttcagagacgacgtaagcgg | Quasar 570 |
| SUN4 | 28 | cagatagcaacatcgttgct | Quasar 570 |
| SUN4 | 29 | ggtaaagagtcacctgcttg | Quasar 570 |
| SUN4 | 30 | aatccttgccaagtgtagta | Quasar 570 |
| SUN4 | 31 | aacgtagtattgcgcggatg | Quasar 570 |
| SUN4 | 32 | tcgacagaaataccggcatt | Quasar 570 |
| SUN4 | 33 | agaaccccaaacacaagcat | Quasar 570 |
| SUN4 | 34 | aattaccgacgcctgaactt | Quasar 570 |
| SUN4 | 35 | agcgccaaagttcaatggag | Quasar 570 |
| SUN4 | 36 | aggcgacaccatcagaagaa | Quasar 570 |
| SUN4 | 37 | caatgcgttaccgttgtttg | Quasar 570 |
| SUN4 | 38 | ctgaagctaccgttttcata | Quasar 570 |
| SUN4 | 39 | aaacttagctttgccagcag | Quasar 570 |
| GENE | Probe # | SEQUENCE | Dye |
| CLB2 | 1 | gcttcttggaaggaaatagc | Quasar 670 |
| CLB2 | 2 | agagagagagggggtaatca | Quasar 670 |
| CLB2 | 3 | tctattgggttggacatcta | Quasar 670 |
| CLB2 | 4 | attctgtgagttttctgtgt | Quasar 670 |
| CLB2 | 5 | taaggccaacctttgtacat | Quasar 670 |
| CLB2 | 6 | gctggattattcgcattact | Quasar 670 |
| CLB2 | 7 | ccttctttactgagtttagt | Quasar 670 |
| CLB2 | 8 | aattgaggaatccgacttcc | Quasar 670 |
| CLB2 | 9 | gatcttgatacgctttctct | Quasar 670 |
| CLB2 | 10 | cattttctttcagggttctt | Quasar 670 |
| CLB2 | 11 | ttgtttttggggagttggat | Quasar 670 |
| CLB2 | 12 | ggtcttggttttctttatca | Quasar 670 |
| CLB2 | 13 | gcaccaaattgctgactact | Quasar 670 |
| CLB2 | 14 | cccctcctttatagaagtta | Quasar 670 |
| CLB2 | 15 | tatatttgcaggcagctcag | Quasar 670 |
| CLB2 | 16 | gctgaggaggattcttgtaa | Quasar 670 |
| CLB2 | 17 | gttatggactacctcagtgt | Quasar 670 |
| CLB2 | 18 | gaagtttttcgttttctacc | Quasar 670 |
| CLB2 | 19 | tgaagttgacttctagctgg | Quasar 670 |
| CLB2 | 20 | gtttcactttcggtatttct | Quasar 670 |
| CLB2 | 21 | gggcagttcttgttcaacaa | Quasar 670 |
| CLB2 | 22 | ccattttcatcgcacacttt | Quasar 670 |
| CLB2 | 23 | ttcccattcatattcttctt | Quasar 670 |
| CLB2 | 24 | ggatcatttacatcttctgc | Quasar 670 |
| CLB2 | 25 | atcattgacgtactcgctga | Quasar 670 |
| CLB2 | 26 | taatgacctctagttggtgg | Quasar 670 |
| CLB2 | 27 | ctccggtaataagccgaatt | Quasar 670 |
| CLB2 | 28 | cctgtccattatgttaatgg | Quasar 670 |
| CLB2 | 29 | atgatgtgccaaccaattgt | Quasar 670 |
| CLB2 | 30 | acctcttcatatttagaggc | Quasar 670 |
| CLB2 | 31 | gtctgtctctgatgcgaaat | Quasar 670 |
| CLB2 | 32 | aaatgaatttctccccttct | Quasar 670 |
| CLB2 | 33 | cggattgggataatttaggt | Quasar 670 |
| CLB2 | 34 | tatcgtagtcatctgctttc | Quasar 670 |
| CLB2 | 35 | atgagggtagtatcccaata | Quasar 670 |
| CLB2 | 36 | cataaacatcgcagctgctg | Quasar 670 |
| CLB2 | 37 | cctttacctaacatttttct | Quasar 670 |
| CLB2 | 38 | tattagatttccatcccatt | Quasar 670 |
| CLB2 | 39 | agttcttctttagtataccc | Quasar 670 |
| CLB2 | 40 | aagcccattggacggaaatt | Quasar 670 |
| CLB2 | 41 | catagccgttttttctaacc | Quasar 670 |
| CLB2 | 42 | ttctcattcatgcaaggtca | Quasar 670 |
| CLB2 | 43 | gcagccatacattttatatg | Quasar 670 |
| CLB2 | 44 | aggcgctaaaattgggtgtg | Quasar 670 |
| CLB2 | 45 | tggtgtggtgttattagtga | Quasar 670 |
| GENE | Probe # | SEQUENCE | Dye |
| ASH1 | 1 | cagcagataatgcatgcagt | Quasar 670 |
| ASH1 | 2 | gctatttgcatgagaatccg | Quasar 670 |
| ASH1 | 3 | atcggttgtgatattgttcc | Quasar 670 |
| ASH1 | 4 | aatacttcctaggacttgct | Quasar 670 |
| ASH1 | 5 | aattggcgacacattgagcg | Quasar 670 |
| ASH1 | 6 | ccaagtatttcgttagccaa | Quasar 670 |
| ASH1 | 7 | gctggtgcagtatttgattt | Quasar 670 |
| ASH1 | 8 | gtaatccatatgatgtggcg | Quasar 670 |
| ASH1 | 9 | aattctggtgaattgcctgg | Quasar 670 |
| ASH1 | 10 | atctgagataagcttgcctt | Quasar 670 |
| ASH1 | 11 | gacccatagttcaatggatt | Quasar 670 |
| ASH1 | 12 | acaatggtagtcgaggttgt | Quasar 670 |
| ASH1 | 13 | aagagacggacgatagcctg | Quasar 670 |
| ASH1 | 14 | agtgatggtaggctttgttg | Quasar 670 |
| ASH1 | 15 | gggtaataattgcagatgcc | Quasar 670 |
| ASH1 | 16 | gtgctgcgttttcttgtaaa | Quasar 670 |
| ASH1 | 17 | acgttttgatgtatcaggga | Quasar 670 |
| ASH1 | 18 | tgggtaagatcagttttcca | Quasar 670 |
| ASH1 | 19 | gtcttgatagtttgtatcct | Quasar 670 |
| ASH1 | 20 | ccacttcttcgcgtatttta | Quasar 670 |
| ASH1 | 21 | gttgggtatacttaatggct | Quasar 670 |
| ASH1 | 22 | caattccttgccgtaattga | Quasar 670 |
| ASH1 | 23 | aacttggacgacctagtcga | Quasar 670 |
| ASH1 | 24 | atgattccttagacggggaa | Quasar 670 |
| ASH1 | 25 | gaggagtaatactccatgca | Quasar 670 |
| ASH1 | 26 | tattggttggtggactcatc | Quasar 670 |
| ASH1 | 27 | aaaaagaggcctctactcct | Quasar 670 |
| ASH1 | 28 | atcgttattgctggatttcc | Quasar 670 |
| ASH1 | 29 | ttccacgtaattgtctttga | Quasar 670 |
| ASH1 | 30 | gcatttggcatgggaaatga | Quasar 670 |
| ASH1 | 31 | actgttcgtctttttgtgac | Quasar 670 |
| ASH1 | 32 | gtctctattcgcaagcaatt | Quasar 670 |
| ASH1 | 33 | tgttcttagatggtttcctt | Quasar 670 |
| ASH1 | 34 | gtcttgcagtgttaccgaaa | Quasar 670 |
| ASH1 | 35 | ggtgaggatgatcttgatct | Quasar 670 |
| ASH1 | 36 | gataattgggtgaccttggg | Quasar 670 |
| ASH1 | 37 | gggagagtcgagagcaaatc | Quasar 670 |
| ASH1 | 38 | atttgatgatcttcgagggc | Quasar 670 |
| ASH1 | 39 | tacttccctttttcgttatg | Quasar 670 |
| ASH1 | 40 | tgtgacgagtgggtgatgag | Quasar 670 |
| ASH1 | 41 | gggcgaatcactcgaatgac | Quasar 670 |
| ASH1 | 42 | gttacatagctgatcttgct | Quasar 670 |
| ASH1 | 43 | aataagcaacggtacccttc | Quasar 670 |
| ASH1 | 44 | ctctactgtctcagttatgt | Quasar 670 |
| ASH1 | 45 | cgcggcgtgtcgaatgaaaa | Quasar 670 |

**Table S2. smFISH probe sequences.** This table shows the sequences of the smFISH probes used for imaging *SRL1*, *SUN4,* *CLB2,* and *ASH1* mRNA in this study. Probes for *SRL1* and *SUN4* were newly designed for this study. Probes for *CLB2* are from Maekiniemi et al. (2025), and for *ASH1* from Tutucci et al. (2018).
